# The genetic history of Mayotte and Madagascar cattle breeds mirrors the complex pattern of human exchanges in Western Indian Ocean

**DOI:** 10.1101/2021.10.08.463737

**Authors:** Jessica Magnier, Tom Druet, Michel Naves, Melissa Ouvrard, Solene Raoul, Jérôme Janelle, Katayoun Moazami-Goudarzi, Matthieu Lesnoff, Emmanuel Tillard, Mathieu Gautier, Laurence Flori

## Abstract

Despite their central economic and cultural role, the origin of cattle populations living in Indian Ocean islands still remains poorly documented. Here, we unravel the demographic and adaptive histories of the extant Zebus from the Mayotte and Madagascar islands using high-density SNP genotyping data. We found that these populations are very closely related and both display a predominant indicine ancestry. They diverged in the 16^th^ century at the arrival of European people who transformed the trade network in the area. Their common ancestral cattle population originates from an admixture between an admixed African zebu population and an Indian zebu that occurred around the 12^th^ century at the time of the earliest contacts between human African populations of the Swahili corridor and Austronesian people from Southeast Asia in Comoros and Madagascar. A steep increase of the estimated population sizes from the beginning of the 16^th^ to the 17^th^ century coincides with the expansion of the cattle trade. By carrying out genome scans for recent selection in the two cattle populations from Mayotte and Madagascar, we identified sets of candidate genes involved in biological functions (cancer, skin structure and UV-protection, nervous system and behavior, organ development, metabolism and immune response) broadly representative of the physiological adaptation to tropical conditions. Overall, the origin of the cattle populations from Western Indian Ocean islands mirrors the complex history of human migrations and trade in this area.

## Introduction

The Indian Ocean has played a prominent role in the human-mediated migration of cattle populations between East-Africa, Middle-East and South-West Asia. However, the origin and genetic diversity of cattle populations living in Indian Ocean islands remain poorly investigated and relatively unclear. Their understanding may provide insights into the recent history of human populations in this area that has long displayed important levels of interaction, trade, migration and domestic species translocation over time (Boivin *et al*. 2013; Beaujard 2019a,b).

Within the large Indian Ocean area, the Comoro Islands (including Ngazidja, Ndzuwani, Mwali and Mayotte) and Madagascar occupy a key position in the maritime trade routes that has linked the East-African coast, Middle East and Asia over the past two millenia. Interestingly, cattle may have been introduced early in this area and have subsequently represented an important domestic species. In both Mayotte and Madagascar where human have permanently settled from the 6th century CE, the most ancient archaeological and skeletal evidence of cattle presence traces back to the 9^th^-10^th^ centuries CE but the quantities of identified cattle bones only increased from 14^th^ - 15^th^ centuries CE (Pauly 2013; Boivin *et al*. 2013). The first Portuguese eyewitnesses reported the presence of cattle on the Western Indian Ocean islands from the 16^th^ century like almost all 17^th^ and 18^th^ century visitors in Comoros (Cheke 2010).

Nowadays, among the Comoros, the cattle population of Mayotte is the best characterized thanks to a recent effort that lead to the official recognition of a local humped breed named ‘Zebu of Mayotte’ (MAY) by the French government for conservation purpose (France 2011; Ouvrard *et al*. 2018). Overall this breed, traditionally and extensively raised in small herds of a handful head, represents 70% of the 20,000 identified individuals identified in the island. Other individuals consist of recently imported individuals from European taurine (EUT) breeds (i.e., Montbeliarde, Jersey, French Brown Swiss, Gasconne) and MAYxEUT admixed individuals. In contrast, nine millions of cattle heads are identified in Madagascar (Ministère de l’Agriculture 2007), including about 85% of Zebu of Madagascar (ZMA) raised in large herds, imported taurine breeds (e.g. Holstein, Norvegian Red), several synthetic admixed breeds with ZMA (such as Renitelo, Manjan’i Boina and Rana) and Baria, a small wild and free-roaming population (Porter 2007, www.fao.org). Both MAY and ZMA populations are considered to be well adapted to their respective islands conditions. In particular, they are generally regarded as more resistant to heat stress and tick-borne diseases than imported European breeds, such as African zebu and N’Dama breeds (Hansen 2004; Mattioli *et al*. 1995; Bock *et al*. 1999; Glass *et al*. 2005).

If the MAY zebu population has not been genetically characterized yet, previous genetic characterization of ZMA populations based on the analysis of a few tens of microsatellite markers or medium density SNP genotyping data suggested an hybrid composition between African taurine (AFT) and indicine (ZEB) ancestries, the latter being predominant (Zafindrajaona and Lauvergne 1993; Hanotte 2002; Gautier *et al*. 2009). However, these analyses remained mostly descriptive and only provide limited insights into the origins of these populations. In particular, from the history of human migration routes, cattle may have been introduced possibly repeatedly with subsequent exchanges between the 8^th^ and the 13^th^ centuries in the Comoro and Madagascar islands from several places including East Africa (as a result of movements of Bantu populations) or Indonesia (with Austronesian navigator populations) (Fuller and Boivin 2009; Fuller *et al*. 2011; Beaujard 2005, 2007, 2011, 2015).

The purpose of our study was to clarify the origin of the local Mayotte and Madagascar cattle breeds to better understand the demography of Western Indian Ocean cattle breeds based on their refined genetic characterization. To that end, we genotyped 32 MAY and 24 ZMA individuals on the bovineHD high density SNP genotyping assay (comprising >770,000 SNPs) together with 113 individuals belonging to the Somba (SOM, n=44) and the Lagune (LAG, n=44) West-African taurine breeds and to the Fulani West-African Zebu (ZFU, n=25). These newly generated data were combined to publicly available bovineHD genotyping data for 363 individuals belonging to 8 other breeds representative of EUT (Angus, Holstein, Jersey and Limousine), another African taurine (NDA), Indian Zebus (Gir and Nelore) and East African Zebus (Bahbahani *et al*. 2017). We first provide a refined analysis of the structuring of genetic diversity among the combined data sets and carried out a detailed inference of the demographic history of the MAY and ZMA populations with respect to the other extant populations by constructing admixture graphs (Patterson *et al*. 2012; Lipson 2020; Gautier *et al*. 2021) and estimating the timing of some admixture events using Linkage-Disequilibrium (LD) information (Loh *et al*. 2013). We further estimated their recent changes in effective population sizes using the recently developed method GONE (Santiago *et al*. 2020) and characterized the levels of genomic inbreeding in MAY and ZMA (Druet and Gautier 2017; Bertrand *et al*. 2019; Druet and Gautier 2021). We finally investigated the patterns of genetic adaptation of MAY and ZMA cattle breeds which recently diverged and live in slightly different tropical island conditions by searching for footprints of positive selection. The identified signals were subjected to a detailed functional analysis to identify putative physiological pathways and their possible underlying selective pressures (see e.g., Flori *et al*. 2019).

## Materials and methods

### Animal sampling and genotyping

For MAY individuals (Zebus from Mayotte), we selected a group of 32 presumably non-related individuals (based on the newly created French National Registration Database) representative of the phenotypic diversity of the local population. They each originated from 32 different farms located in 17 townships spread over the Grande and Petite terres of the Mayotte Island (Figure S1, Table S1). Blood samples were collected during year 2016 from the tail vein of the individuals using 10 ml EDTA vacutainer tubes, strictly following the recommendations of the directive 2010/63/EU for animal care. Genomic DNA was further extracted using the Wizard Genomic DNA purification kit (Promega, France) and stored at −20°C. MAY DNA samples were genotyped (Table S1) on the Illumina BovineHD genotyping beadchip at Labogena plateform (Jouy-en-Josas) using standard procedures (www.illumina.com) together with 24 Zebus of Madagascar (ZMA) and 113 West-African cattle DNA samples (i.e. 44 Lagune, 44 Somba and 25 Zebus Fulani), sampled in the 1990’s and previously genetically characterized on the Illumina BovineSNP50 beadchip (Gautier *et al*. 2009). The genotyping data were then added to the WIDDE database and combined with other publicly available ones, using WIDDE utilities (Sempéré *et al*. 2015).

To obtain the position of the 777,962 SNPs included in the BovineHD genotyping assay onto the latest ARS-UCD1.2 (aka bosTau9) bovine genome assembly (Rosen *et al*. 2020), the SNP sequences were realigned onto this assembly using pblat (Wang and Kong 2019) run with options -out=pslx -minIdentity=98 -threads=4. The resulting pslx file was parsed with a custom awk script to obtain SNP positions which were unambiguous for 721,583 SNPs (92.8%). We discarded the 15,533 (2.0%) SNPs that mapped to more than 9 different positions but kept the remaining 40,846 SNPs with 2 (n=19,907) to 9 different alignments assigning to them the position given by the one with the highest score.

The complete genotyping dataset consisted of 532 animals including 363 individuals from 8 other bovine populations (Bahbahani *et al*. 2017, www.illumina.com) representative of the bovine worldwide genetic diversity (Table 1). The minimal individual genotyping call rate was set to 90% and the minimal SNP genotyping call rate to 90% over all populations and 75% within each population (i.e. SNPs genotyped for less than 75% of the animals from at least one population were discarded). SNPs with a MAF*<*0.01 or departing from Hardy-Weinberg equilibrium expectation (exact test P-value*<* 10^−3^ in at least one breed) were also discarded. A total of 680,338 SNPs distributed throughout the 29 autosomes of the bosTau9 bovine genome assembly passed finally all our filtering criteria.

**Table 1.** Sample description and origin of the Illumina BovineHD chip genotyping data (^⋆^ before filtering).

| Code | Population name | Land of origin | Sampling Year | Nb | Reference |
| --- | --- | --- | --- | --- | --- |
| ANG | Angus | Scotland | n.a. | 42 | Illumina (Sempéré et al. 2015) |
| EAZ | East African Shorthorn Zebu | Kenya | n.a. | 92 | Bahbahani et al. (2017) |
| GIR | Gir | India | n.a. | 27 | Illumina (Sempéré et al. 2015) |
| HOL | Holstein | Netherlands | n.a. | 60 | Illumina (Sempéré et al. 2015) |
| JER | Jersey | Jersey | n.a. | 38 | Illumina (Sempéré et al. 2015) |
| LAG | Lagune | Benin | 1996 | 44 | This study |
| LMS | Limousine | France | n.a. | 50 | Illumina (Sempéré et al. 2015) |
| MAY | Zebu from Mayotte | Mayotte | 2016 | 30 (32)* | This study |
| NDA | N'Dama | Guinea | n.a. | 23 | Illumina (Sempéré et al. 2015) |
| NEL | Nelore | India | n.a. | 31 | Illumina (Sempéré et al. 2015) |
| SOM | Somba | Togo | 1996 | 44 | This study |
| ZFU | Zebu Fulani | Benin | 1996 | 24 (25)* | This study |
| ZMA | Zebu from Madagascar | Madagascar | 1991 | 23 (24)* | This study |

### Inference of the population demographic history

#### Characterization of the structuring of genetic diversity

Principal Component Analysis (PCA) based on individual SNP genotyping data was performed with *smartpca* (Patterson *et al*. 2006) and visualized with the R package ggplot2 (Wickham 2016). Unsupervised genotype-based hierarchical clustering of the individual animal samples was carried out using the maximum-likelihood method implemented in ADMIXTURE 1.06 (Alexander *et al*. 2009). Results were visualized with custom functions in R environment (www.r-project.org). Finally, the overall and pairwise-population *F*_*ST*_ were computed with version 2.0 of the R package poolfstat (Gautier *et al*. 2021) using the computeFST ran with default settings (i.e., method=Anova) and option nsnp.per.bjack.block=5000 to estimate standard errors (and 95% CI as ±1.96 s.e.) with block-jackknife.

#### f-statistics based demographic inference

*f*-statistics based demographic inference (Patterson *et al*. 2012) were carried out with the new functionalities of the version 2.0 of the R package poolfstat (Gautier *et al*. 2021). We used the compute.fstats function to estimate the different *f*-statistics (including *F*_3_ and *F*_4_ for all the population triplets and quadruplets respectively) and within-population heterozygosities. As for *F*_*ST*_, standard-errors of the estimated statistics (and their corresponding Z-scores for *f*_3_ and *f*_4_) were estimated using block-jackknife defining blocks of 5,000 consecutive SNPs (i.e., option nsnp.per.bjack.block=5000). In addition, to mitigate SNP ascertainment bias by favoring SNPs of most remote ancestry (e.g., discard SNPs of exclusive European ancestry), we only kept for *f*-statistics based demographic inference the 497,949 polymorphic SNPs that were polymorphic (MAF>0.05) in both ZEB (GIR and NEL combined) and in AFT (NDA, SOM and LAG combined) populations. Following Patterson *et al*. (2012) (see also Lipson 2020), we then carried out formal tests of population admixture using the estimated *f*_3_ statistics. A negative Z-score (*Z <* − 1.65 at the 95% significance threshold) associated to an *f*_3_ for a given population triplet A;B,C showing that the target population A is admixed between two source populations each related to B and C. To further provide insights into the origins of MAY and ZMA, we build an admixture graph with poolfstat utilities (Gautier *et al*. 2021). Briefly, we first build a scaffold tree of presumably unadmixed populations (as suggested by both exploratory analyses and *f*_3_ and *f*_4_ based tests) consisting of two AFT breeds (NDA and LAG), two ZEB breeds (GIR and NEL) and one EUT breed (HOL) with the rooted.njtree.builder function. We further relied on the *graph*.*builder* function (ran with default options) to jointly include EAZ, MAY and ZMA that showed clear evidence for admixture on the graph considering all the six possible orders of inclusion. The fit of the best fitting graph (displaying a BIC more than 8 units lower than all the other graphs explored in the graph building process) was further validated with the *compare*.*fitted*.*fstats* function that allows to compare to which extent the estimated *f*-statistics depart from their predicted values based on the fitted admixture graph parameters via a Z-score (Patterson *et al*. 2012; Lipson 2020; Gautier *et al*. 2021).

#### Estimation ot the timing of admixture events

We estimated the timing of admixture events (in generations) with the program mALDER (Pickrell *et al*. 2014) that implements a modified version of ALDER method originally described by Loh *et al*. (2013). This approach relies on the modelling of the exponential decay of admixture-induced LD in a target admixed population (here based on two-reference weighted LD curves, i.e., using a LD measure weighted by allele frequencies in two source population proxies) as a function of genetic distance. Genetic distances between pairs of SNPs were derived from physical distances assuming a cM to Mb ratio of 1 (Kadri *et al*. 2016). A six year generation time was assumed to convert the timing from generations to years (Gautier *et al*. 2007; Keightley and Eyre-Walker 2000).

#### Inference of the recent population size histories

Historical effective population sizes (*N*_*e*_) were inferred for the MAY and ZMA breeds with the program GONE that implements an approach recently developed by Santiago *et al*. (2020) to fit the observed spectrum of LD of pairs of loci over a wide range of recombination rates (which we derived from physical map distances assuming a cM to Mb ratio equal to 1, see above). In practice, we adopted a block-jackknife approach to estimate confidence intervals for the inferred *N*_*e*_ trajectories by first identifying 55 non-overlapping blocks of 10,000 consecutive SNPs (block size ranging from 32.8 to 40.2 Mb) out of the 680,338 genotyped ones. We then analyzed 55 different data sets of 670,338 SNPs that were each formed by removing from the original data sets one block of 10,000 SNPs. The resulting 55 inferred *N*_*e*_ trajectories were then summarized by computing a mean trajectory and a 95% confidence envelope defined by the 2.5 and 97.5 percentiles from the 55 estimated *N*_*e*_ at each time point.

#### Age-based partitioning of individual inbreeding in MAY and ZMA breeds

The characterization of individual levels of inbreeding at a global and local scale in MAY and ZMA breeds was performed with the model-based approach implemented in the RZooRoH package (Bertrand *et al*. 2019; Druet and Gautier 2017, 2021). This method models each individual genomes as a mosaic of Homozygous-by-Descent (HBD) and non-HBD segments using a multiple HBD-classes Hidden Markov Model (Druet and Gautier 2017). Each HBD class is specified by a rate *R*_*k*_ related to the expected length (equal to 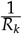 Morgans) of the associated HBD segments and that is approximately equal to twice the number of generations to the common ancestor that transmitted the DNA segment. Given the density of the HD chip, we considered a model with 11 HBD classes (with *R*_*k*_ = 2^*k*^ for *k* = 1, …, 11) and one non-HBD class (with rate equal to *R*_11_) allowing to capture the contribution to the overall individual inbreeding levels from each age-based classes of ancestors (living up to 1,024 generations in the past).

### Whole-genome scan for footprints of selection

#### Computation of iHS and Rsb statistics

The genome-wide scan for footprints of positive selection within and between MAY and ZMA breeds was performed using extended haplotype homozygosity (EHH)-based tests. To that end we first jointly phased the genotyping data for MAY, ZMA, EAZ, GIR and NEL individuals with version 1.4 of the fastPHASE software (Scheet and Stephens 2006) for each chromosome in turn using population label information and options -Ku40 -Kl10 -Ki10. Genetic distances between consecutive SNPs were obtained from physical distances assuming a cM to Mb ratio equal to 1 (see above). Based on haplotype information, we further computed the *iHS* (Voight *et al*. 2006) and *Rsb* (Tang *et al*. 2007) statistics using the version 3.1.2 of the R package rehh (Gautier *et al*. 2017; Klassmann and Gautier 2020). These statistics are designed to detect regions with high level of haplotype homozygosity over an unexpected long distance (relative to neutral expectations), either within population (*iHS*) or across populations (*Rsb*). Note that for the computation of *iHS* within MAY and ZMA populations, we chose to not polarize the SNP alleles (i.e., scan_hh function was run with option polarized=FALSE) as discussed in section 7.6 of the online rehh package vignette (https://cran.r-project.org/web/packages/rehh/vignettes/rehh.html). The different SNP *iHS* and *Rsb* statistics were further transformed into *p*_*iHS*_ = − log_10_ (1 − 2 | Φ(*iHS*) − 0.5 |) and *p*_*Rsb*_ = − log_10_ (Φ(*Rsb*)) where Φ(*x*) represents the Gaussian cumulative function. Assuming *iHS* and *Rsb* scores are normally distributed under neutrality, *p*_*iHS*_ and *p*_*Rsb*_ might thus be a twosided − log_10_ (P-value) associated with the neutral hypothesis.

#### Identification of candidate genes

We used the approach previously described in Flori *et al*. (2019) to identify candidate genes under positive selection from *iHS* and *Rsb* estimates. Briefly, all the genotyped SNPs were annotated using as a gene set reference a list of 14,562 RefSeqGenes anchored on the Btau9 bovine genome assembly (refGene.txt.gz, 2019-06-07; https://hgdownload.soe.ucsc.edu/goldenPath/bosTau9/database/). A SNP was then considered as representative of one of these RefSeq genes if localized within its boundary positions extended by 15 kb upstream and downstream to account for persistence of LD (e.g., Gautier *et al*. 2007). Among the 680,338 SNP of HD-dataset participating to the analysis, 293,527 SNPs mapped to 13,696 different RefSeq Genes, corresponding to 13,536 gene symbols. On average, each SNP mapped within 1.2 RefSeq genes (from 1 to 34, median= 1) and each RefSeq gene was represented by 21 SNPs (from 1 to 641, median=13). For a given statistic, genes with at least 2 representative SNPs with −log_10_ (P-value) *>* 4 were considered as candidate genes.

#### Functional annotation of candidate genes under selection

Following the approach outlined in Flori *et al*. (2019), candidate genes were functionally annotated and submitted to gene network analysis using Ingenuity Pathway Analysis software (IPA, QIAGEN Inc., https://www.qiagenbioinformatics.com/products/ingenuity-pathway-analysis). Among the 13,535 genes symbols taken into account in the analysis, 12,324 were mapped in the Ingenuity Pathway Knowledge Base (IPKB) and were considered as the reference data set. Among the candidate genes identified for MAY (N=27) and ZMA (N=47) breeds, 24 and 45 were respectively mapped to IPKB. The top significant functions and diseases (P-value*<* 0.05) were obtained by comparing functions associated with the candidate genes under selection against functions associated with all genes in the reference set, using the right-tailed Fisher exact test. In the network analysis, a score S was computed for each network that contained at most 140 molecules (including candidate genes under selection) based on a right-tailed Fisher exact test for the over-representation of candidate genes under selection (*S* = − log_10_ (P-value)). A network was considered as significant when *S >* 3. The top significant functions and diseases associated with significant networks were also reported.

## Results

### Genetic relationships of Mayotte and Madagascar cattle breeds with other world cattle breeds

We first carried out an individual-based PCA of the SNP genotyping dataset consisting of 680,338 autosomal SNPs genotyped on individuals from MAY (n=32) and ZMA (n=23) populations together with 473 individuals belonging to 11 cattle breeds representative of the bovine genetic diversity (Table 1). The first factorial plan of the PCA is represented in Figure 1a. The first two components accounting for 21.09% and 7.85% of the total variation respectively. In agreement with previous studies (e.g., Gautier *et al*. 2010), the first two PCs revealed a clear structuring of individual genetic diversity according to their population of origins and highlighted a triangle-like 2-dimensional global structure of the cattle populations with apexes corresponding to three main groups: i) European taurines (EUT) represented by ANG, HOL, JER and LMS; ii) African taurines (AFT) represented by LAG, NDA and SOM; and iii) zebus (ZEB) represented by NEL and GIR. Similarly, as previously observed with data from medium density SNP genotyping data (on the same sample), ZMA and West-African hybrids (ZFU and EAZ) were located at intermediate positions on the AFT/ZEB segment, the ZMA individuals being closer to the ZEB apex (Gautier *et al*. 2010). Finally, the newly characterized MAY individuals appeared mostly confounded with ZMA individuals which is consistent with their close geographical proximity. Unsupervised hierarchical clustering analysis of the individuals using K=3 predefined clusters provided a complementary view in close agreement with PCA results while allowing to provide first estimates of ancestry proportions (Figure 1b). Roughly interpreting the blue, red and green cluster in Figure 1b as representative of EUT, ZEB and AFT ancestries respectively suggested that ZMA and MAY individuals had similarly high levels of ZEB ancestry (0.83 ± 0.02 and 0.82 ± 0.01 on average, respectively), a weaker level of AFT ancestry (0.16 ± 0.01), and almost no detectable EUT ancestry except for three individuals (2 MAY and 1 ZMA) with more than 5% of EUT ancestry, which were discarded from further analyses.

**Figure 1.**
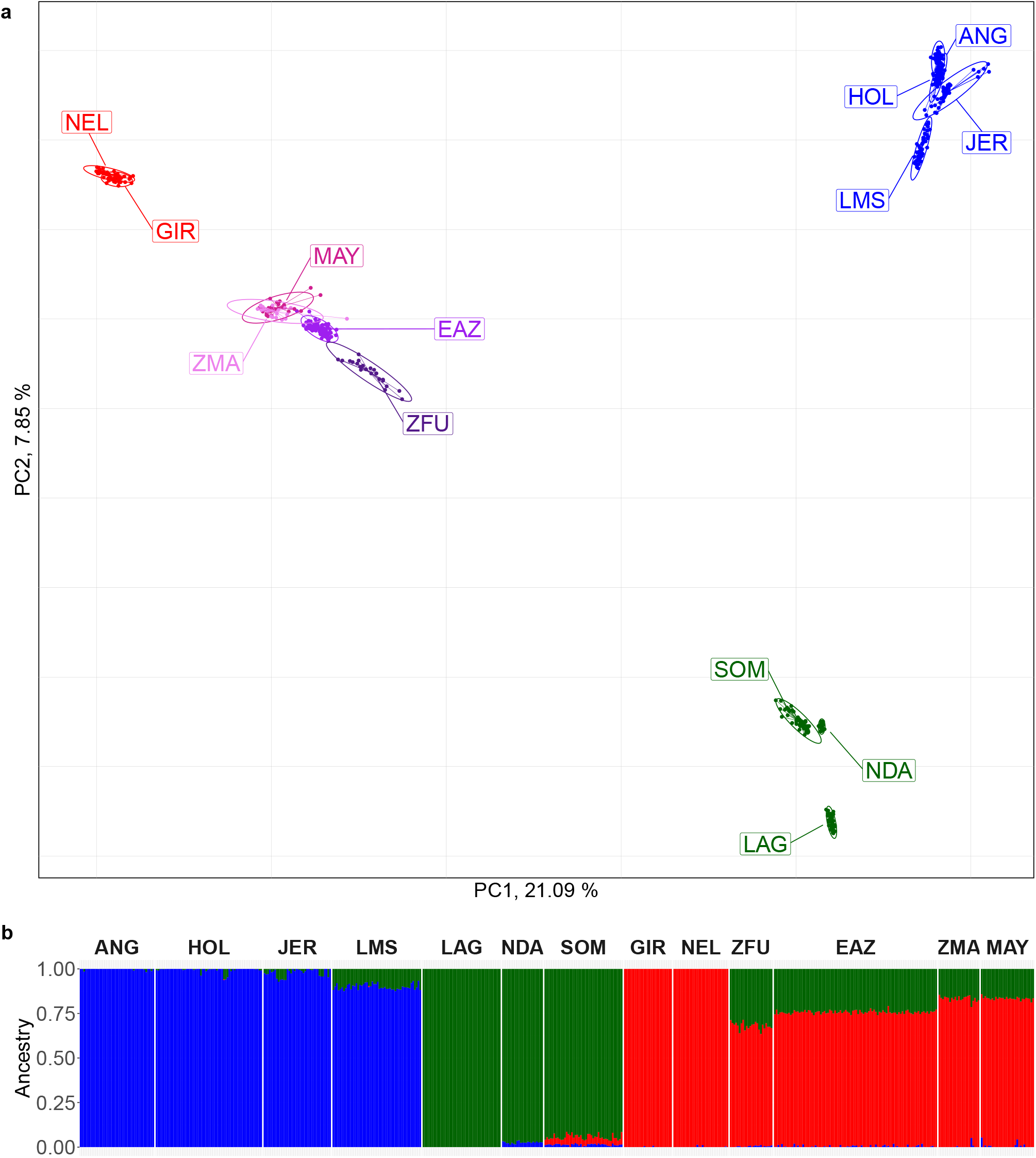
Results of the Principal Component Analysis (PCA) and unsupervised hierarchical clustering including HD genotyping data (528 individuals from 13 populations genotyped for 680,338 SNPs). **a** PCA results. The individuals are plotted on the first two principal components according to their coordinates. Ellipses characterize the dispersion of each population around its centre of gravity. MAY and ZMA individuals are plotted in dark-pink and pink and EAZ and ZFU individuals in purple and dark purple, respectively. EUT, ZEB and AFT individuals are plotted in blue, red and green. **b** Unsupervised hierarchical clustering results with K=3 predefined clusters. For each individual, the proportions of each cluster (y axis) which were interpreted as representative of EUT, AFT and ZEB ancestries are plotted in blue, green and red, respectively.

The different analyses performed at an individual scale showed that the partitioning of cattle into distinct populations (and breeds) is relevant to study their genetic history at the population level (e.g., Gautier *et al*. 2010). Accordingly, the overall *F*_*ST*_ among the 13 populations was found equal to 0.271 (95% CI, [0.262; 0.279]). Population pairwise *F*_*ST*_ ranged from 2.40% (± 0.01%) for the ZMA/MAY pair to 51.1% (± 0.06%) for the LAG/NEL pair (Figure S2) confirming the close relatedness of this two populations. Among the 11 other populations, EAZ was found the most closely related to both MAY and ZMA (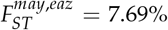 and 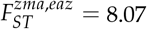) followed by ZFU and the two ZEB populations (GIR and NEL).

### Inferring the history of Mayotte and Madagascar cattle populations

#### f_3_-based tests shows clear evidence for admixture in the history of both MAY and ZMA

Out of the 66 (= 12) possible *f*_3_-based tests when considering either MAY or ZMA as a target population, six were significantly negative (Z-score*<* − 2.33 at the 99% significance threshold) providing strong evidence for admixture events in their history. They all involved the same pair of source proxies that consisted of one AFT population (LAG, NDA or SOM) and one ZEB population (NEL or GIR), the most significant signal being observed with the LAG/NEL pair for ZMA (*f*_3_ (ZMA;LAG,NEL) associated Z-score equal to −3.92) and the NDA/NEL pair for MAY (*f*_3_ (MAY;NDA,NEL) associated Z-score equal to −6.38). Among the 11 other populations, EAZ, ZFU and SOM also showed clear evidence for admixture with 14, 14 and 6 *f*_3_-based tests with a significantly negative Z-score (at the 99% significance threshold), respectively. For each of these three target populations, the lowest Z-score was always associated with the LAG/NEL pair of source proxies (*f*_3_ (EAZ;LAG,NEL), *f*_3_ (ZFU;LAG,NEL) and *f*_3_ (SOM;LAG,NEL) associated Z-scores being equal to −36.5, −33.6 and −9.13 respectively).

#### admixture graph construction

To infer the history of MAY and ZMA ancestry, we constructed an admixture graph relating these two populations with two out of the three AFT breeds (LAG and NDA, discarding SOM as it displayed strong signal of presumably recent admixture with ZEB ancestry, see above), the two ZEB breeds (GIR and NEL), one (HOL) out of four EUT breeds and the EAZ African hybrid. We did not seek to include ZFU as it was more remotely related to East-African populations with a recent origin not directly informative for the MAY and ZMA history (Flori *et al*. 2014). The inferred best fitting admixture graph is represented in Figure 2. It shows that MAY and ZMA breeds both derived from an admixed population we referred to Indian Ocean Zebu in Figure 2, although it should be noticed that we have no clear evidence about the actual geographic origin of this ancestral population. This Indian Ocean Zebu had an admixed origin with a predominant ancestry (88.6%) from an admixed Zebu, likely of African origin since the most closely related to extant EAZ, and the remaining ancestry (11.4%) originating from a presumably Indian Zebu related to extant GIR. The former African Zebu was itself admixed with a Zebu related to extant NEL and a Taurine related to African taurine ancestries (in 68.4% vs. 31.4% respective proportions). Note that these inferred origins lead to overall ZEB (resp. AFT) related ancestries for MAY and ZMA similar to the one obtained in Figure 1.

**Figure 2.**
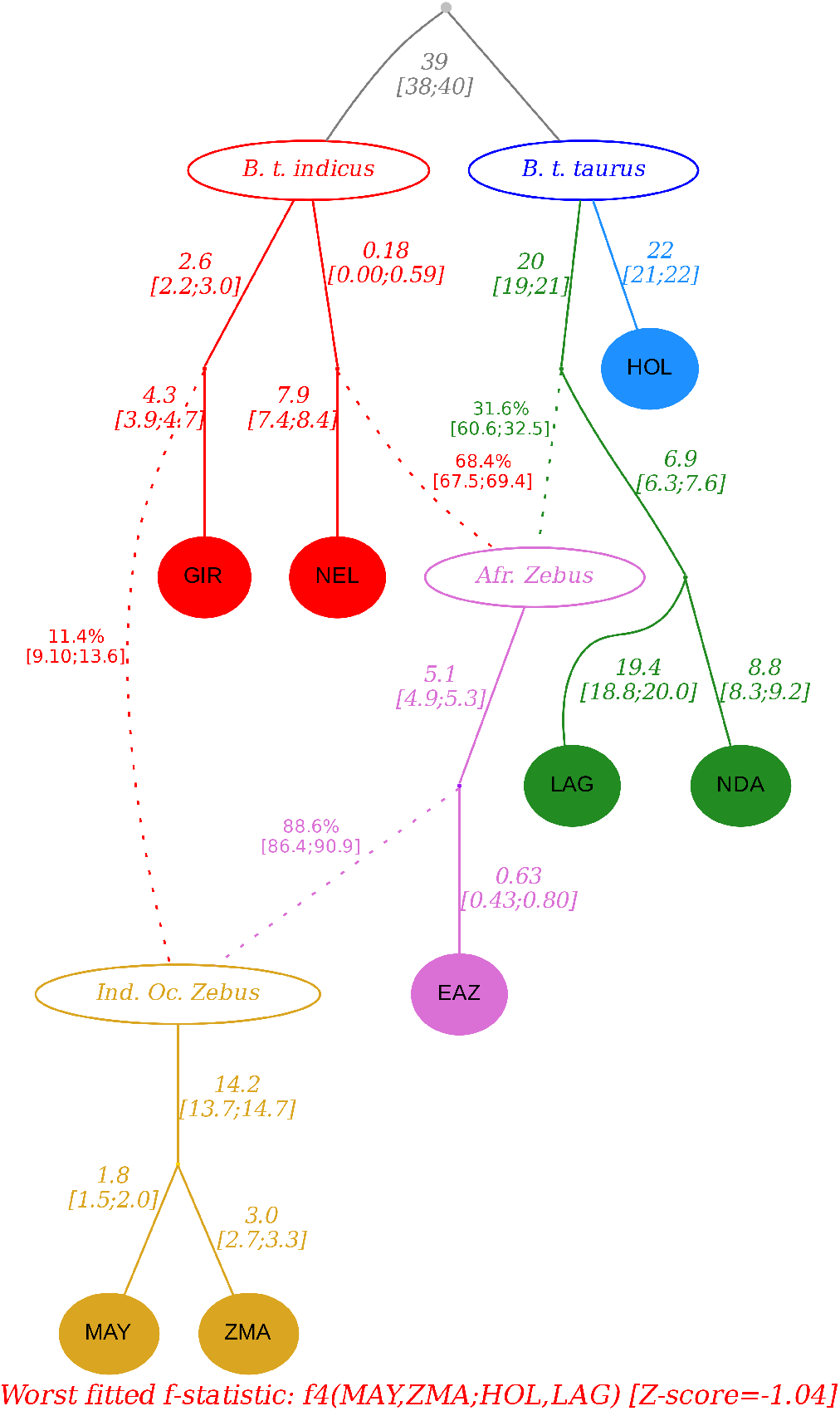
Inferred admixture graph connecting cattle breeds from Mayotte and Madagascar (MAY and ZMA, in yellow) with two Indian indicine breeds (GIR and NEL in red), one African zebu breed (EAZ in purple), two African taurine breeds (LAG and NDA in green) and one European taurine breed (HOL in blue). Admixture events are shown by dotted arrows. Estimates of branch lenghts (×10^3^, in drift units of 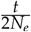) and admixture rates are indicated next to the corresponding edges.The Z-score of the worst fitted *f*-statistics *f4*(MAY,ZMA;HOL,LAG) is equal to −1.04.

#### timing of admixture events

We estimated the timing of the two aforementioned admixture event 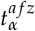 (leading to African Zebus, closely related to EAZ) and 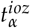 (leading to Indian Ocean Zebus, the direct ancestor of MAY and ZMA) with the program mALDER Pickrell *et al*. (2014). Based on the inferred admixture graph (Figure 2), we estimated 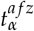 by using EAZ as a population target and a pair of source population proxies consisting of one AFT (either NDA or LAG) and one ZEB (either GIR or NEL). In agreement with admixture graph fitting, the highest amplitude (i.e., *y*-intercept) estimate of the fitted weighted LD curve was obtained with NDA and NEL as source proxies which suggests that these populations are the closest (among the sampled ones) to the actual source population (Loh *et al*. 2013). The corresponding estimated timing for the admixture events leading to African Zebus was found equal to 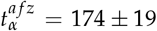 generations (i.e., 1, 044 ± 114 years). This admixture events thus traces back to the 10^th^ century CE (ca. year 950±150). We similarly estimated 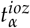 by considering either MAY or ZMA as a population target and a pair of source population proxies consisting of EAZ and one ZEB (either GIR or NEL). For both target populations, the amplitude was the highest (although only slightly) with GIR as the ZEB represent. The estimated timing for the admixture events leading to Indian Ocean Zebus was found equal to 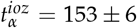 and 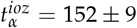 generations with MAY and ZMA as a target population respectively. The (slightly) lower latter estimate being consistent with the difference in sampling time (ca. 4 generations) between MAY and ZMA samples. This second admixture events thus traces back to the early 12^th^ century CE (ca. year 1,100±40).

#### recent population size history of MAY and ZMA

Figure 3 plots the recent effective population size histories of MAY and ZMA as inferred with GONE (Santiago *et al*. 2020). The two trajectories were found similar before the beginning of the 16^th^ century (i.e., ca. 80 generations before MAY sampling time) when the two populations likely started to diverge. From that time onward, the two populations experienced a rapid growth in a few generations toward a mostly constant size (approximately twice higher for ZMA and MAY) followed by a new moderate increase from the beginning of the 20th century. As expected from the large differences in census cattle population sizes between Mayotte and Madagas-car islands, the current estimated *N*_*e*_ was about twice lower in MAY compared to ZMA. More precisely, the harmonic mean of the (mean) historical *N*_*e*_ from generation 80 (76 for ZMA) onward was found equal to 2,160 and 1,045 for ZMA and MAY respectively. Surprisingly, a close inspection of the inferred admixed graph in Figure 2 seems to contradict these large differences in *N*_*e*_ since the estimated (leave) branch lengths was about twice higher for ZMA compared to MAY. This thus rather suggested increased drift in ZMA since these estimates (on a diffusion timescale) might be interpreted under a pure-drift model of divergence as inversely proportional to *N*_*e*_ (i.e., equal to 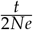 where *t* is the divergence time in generation). Population-specific *F*_*ST*_ estimates for MAY and ZMA essentially confirmed this trend (1.59% and 3.15% respectively, Figure S2). Conversely, when considering only the 518,315 SNPs with a MAF*>*1% in MAY and ZMA (computed over MAY and ZMA combined individuals), the estimated heterozygosity was slightly higher in MAY (33.2%) than in ZMA (32.7%). As detailed in Figure S3, these apparent discrepancies may actually be explained by some residual asymmetric gene flow between MAY and ZMA (or from a common gene pool related to some other continental population, e.g., from the East-African coast) after these two population split. More specifically, very similar level of differentiation and heterozygosities among MAY and ZMA could be obtained with data simulated with msprime (Kelleher *et al*. 2016) under a simplified scenario assuming MAY and ZMA split 80 generations ago and maintained a constant *N*_*e*_ (equal to the one estimated above) with MAY receiving twice as many migrants as ZMA from a common gene pool (*m*_→*MAY*_ = 4 × 10^−3^ vs *m*_→*ZMA*_ = 2 × 10^−3^) (Figure S3).

**Figure 3.**
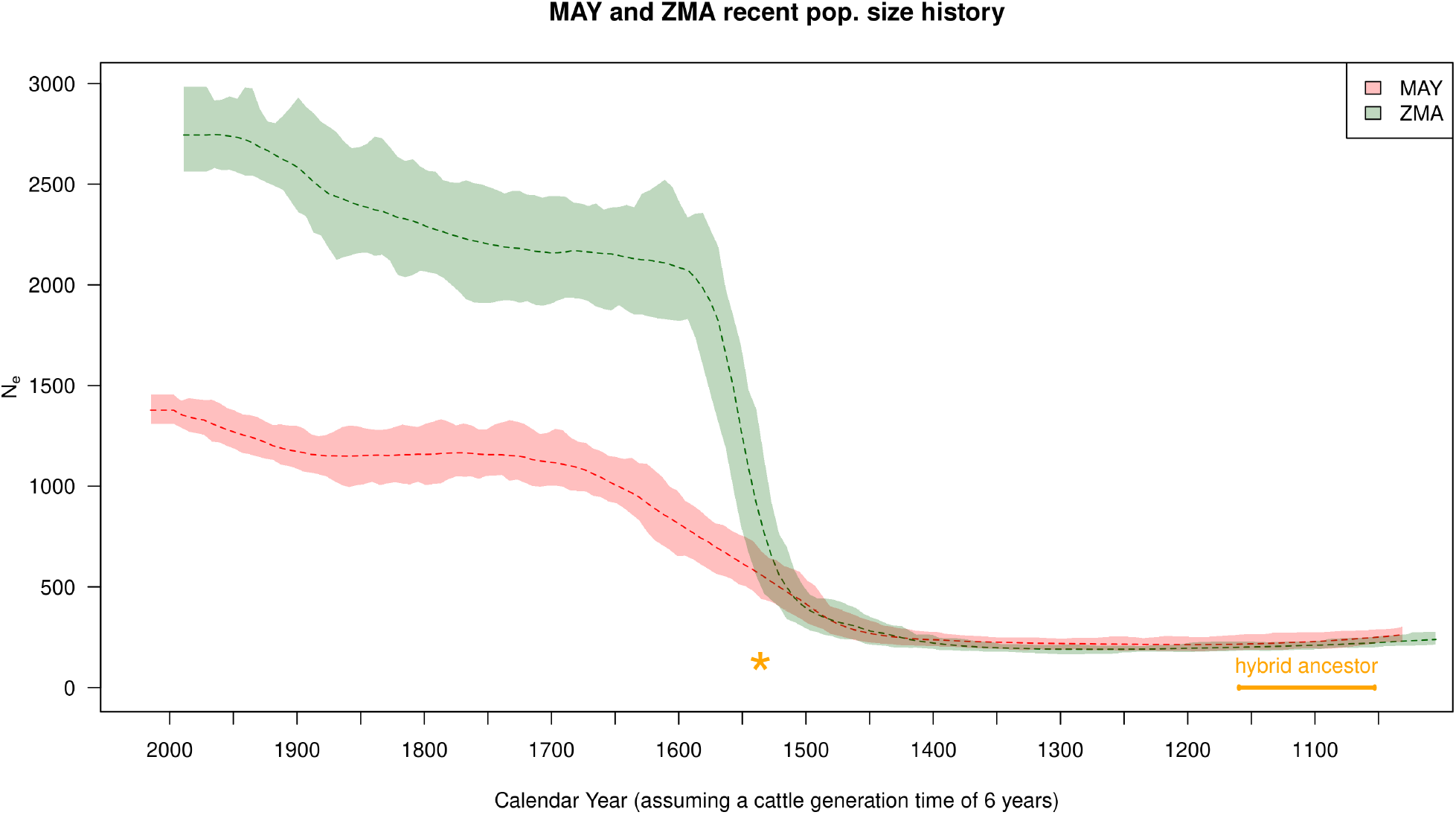
Population size history (*N*_*e*_) of MAY and ZMA population estimated with GONE (Santiago *et al*. 2020). The average *N*_*e*_ trajectories (dashed line) and 95% confidence envelope estimated from block-jackknife samples are plotted in green for ZMA and red for MAY. The time scale was transformed into calendar assuming a 6-year generation time for cattle and accounting for the difference in sampling time between ZMA (1990) and MAY (2016). The estimated timing of the admixture event 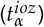 that led to the common hybrid ancestor (named Indian Ocean Zebus in the main text) of MAY and ZMA is given in orange. The orange asterisk gives the likely splitting time (beginning of the 16th century, i.e., *t*_*S*_ ≃ 80 generations before MAY sampling time) between MAY and ZMA roughly estimated from the separation of the two trajectories.

#### MAY and ZMA individuals levels of inbreeding

In order to characterize individual levels of recent inbreeding at both a global (genome-wide) and local (locus-specific) scale, we estimated the individual inbreeding coefficients in MAY and ZMA populations by applying the method implemented in RZooRoH software (Druet and Gautier 2017; Bertrand *et al*. 2019). The distribution of the genome-wide coefficients of individual inbreeding estimated with the R package RZooRoH are detailed in Figure 4a for the MAY and ZMA populations together with their age-based partitioning to assess the contribution of various class of ancestors (Figures 4b, 4c, Figure S4). Overall, the average individual inbreeding levels were found similar and equal to 0.308 ± 0.06 and 0.3136 ± 0.03 in MAY and ZMA respectively, MAY and ZMA individuals all displaying at least 16% and 25% of their genome in autozygous (HBD) segments. However, in agreement with the inferred trajectories of historical *N*_*e*_ (Figure 3, most HBD segments were concentrated in ancestor age classes associated with *R*_*k*_ equal to 128 and 256 (i.e. approximately 64 and 128 generations before sampling thus spanning the split time period between MAY and ZMA). Hence, most of the estimated inbreeding level originated from the foundation of the two populations. Consistently, only a few individuals displayed a high proportion of HBD segments associated with the most recent common ancestors, two generations ago. More precisely, two MAY individuals showed more than 49% of their genome in HBD classes and two ZMA individuals showed more than 40% of their genome classified has HBD.

**Figure 4.**
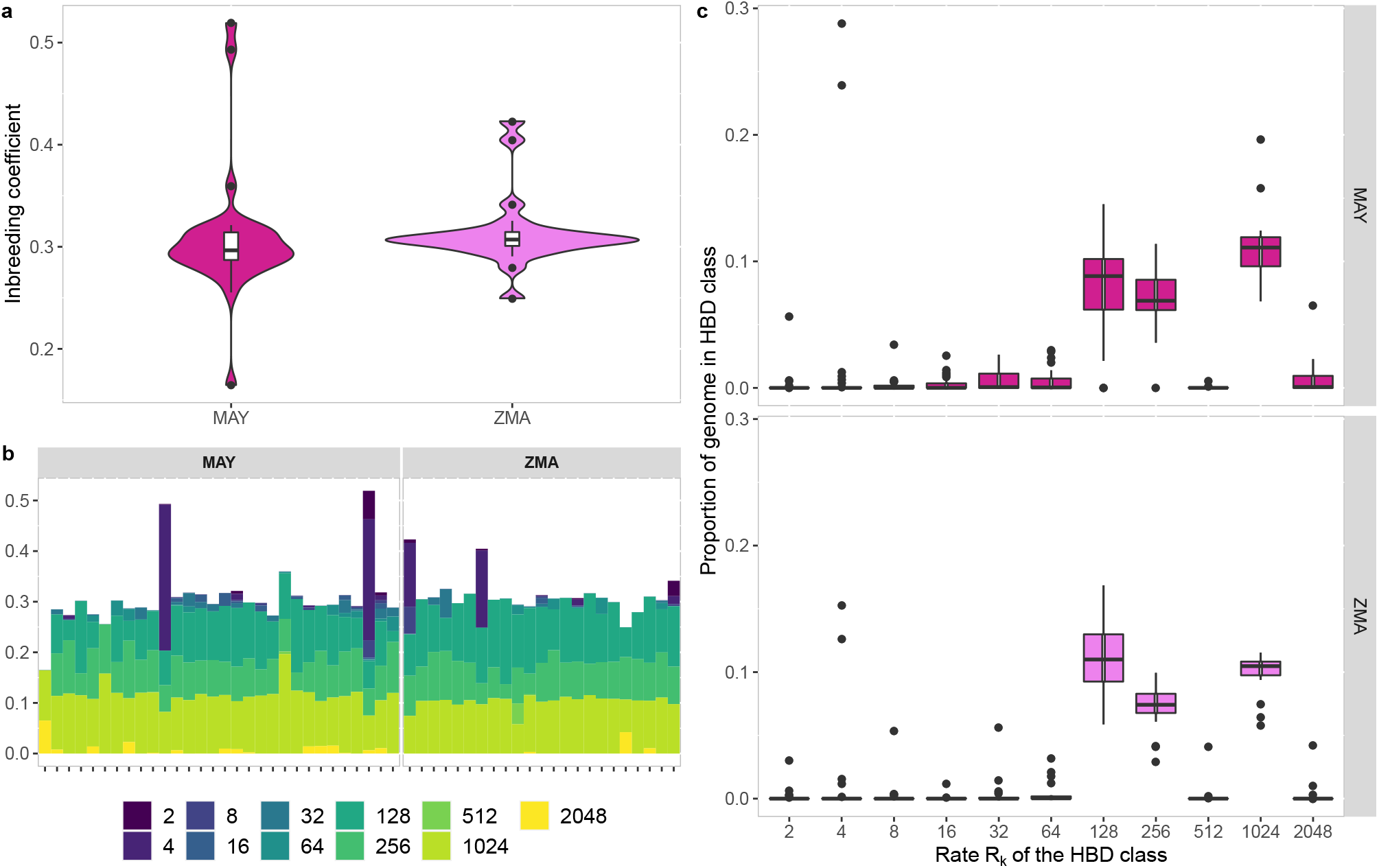
Characterization of individual inbreeding levels in Mayotte and Madagascar breeds. **a** Violin plots representing the distribution of inbreeding coefficients for the MAY (N=30) and ZMA (N=23) breeds, coloured pink and purple, respectively. **b** Partitioning of individual genomes in different HBD classes. Each bar represent an individual and its total height the overall level of inbreeding. The height of the different stacks, which appear in different colors, represents the proportion of the genome associated with each HBD class, defined by their rate *R*_*k*_. **c** Average proportion of individual genomes associated with different HBD classes for MAY (pink) and ZMA (purple) populations. Individual proportions of the genome associated with a specific HBD class are obtained by averaging the corresponding HBD-classes probabilities over all marker positions.

### Identification of selection footprints in the MAY and ZMA genomes

The genome of ZMA and MAY individuals was scanned for footprints of selection with the package *rehh* 3.1.2 (Gautier *et al*. 2017), following the approach described in Flori *et al*. (2019) to identify directly candidate genes. According to our criteria (see Material and Methods), the computation of the *iHS* statistic for each SNP over the whole MAY and ZMA genomes detected two (i.e. GRIK2 and RBM47) and 12 (i.e. AASS, ACTR6, ATP10D, COL15A1, DPP6, DPYSL5, EXOC4, ITGA7, MAPRE3, SLC17A8, TNFSF11, ZCCHC7) genes under selection located on two (i.e. BTA6 and 9) and six (i.e. BTA4, 5, 6, 8, 11 and 12) chromosomes, respectively (see details in Table S2; Figures S5 and S6), with no overlap between these two lists of genes. By computing different *Rsb* statistics comparing the local extend of haplotype homozygosities between MAY and ZMA, and between these two breeds and each of their different main ancestries (i.e. AFZ and ZEB), 3, 16, 12, 18, 11 and 11 candidate genes were identified with the *Rsb*_*MAY*/*ZMA*_, *Rsb*_*MAY*/*EAZ*_, *Rsb*_*MAY*/*ZEB*_, *Rsb*_*ZMA*/*MAY*_, *Rsb*_*ZMA*/*EAZ*_ and *Rsb*_*ZMA*/*ZEB*_ tests, respectively (Table S2, Figure S5 and S6). Most of the candidate genes identified by the *Rsb* involving MAY or ZMA populations were different (74 and 85%, respectively). However seven candidates genes (i.e. GRIK2, HMGSC2, LPCAT2, MIR2285BM, PHGDH, TRIM10 and TRIM15) were identified under selection in both breeds by some of these tests. Specifically, GRIK2, located on BTA9 was found under selection by five tests (*iHS*_*MAY*_, *Rsb*_*MAY*/*EAZ*_, *Rsb*_*MAY*/*ZEB*_ and *Rsb*_*ZMA*/*EAZ*_ and *Rsb*_*ZMA*/*ZEB*_ tests), HMGSC2 and PHGDH located on BTA3 and LPCAT2 located on BTA18 by *Rsb*_*MAY*/*ZEB*_ and *Rsb*_*ZMA*/*ZEB*_ tests, MIR2285BM located on BTA16 by *Rsb*_*MAY*/*ZEB*_ and *Rsb*_*ZMA*/*EAZ*_ tests and TRIM10 and 15 located on BTA23 by *Rsb*_*MAY*/*EAZ*_ and *Rsb*_*ZMA*/*EAZ*_ tests. *Rsb*_*MAY*/*ZMA*_ test detected three (i.e. LRRK2, GSTCD and GPC5) and 18 genes (i.e. ATP2C1, SCN2A, STPG1, KCND2, ANKS1B, NR1H4, MIR2434, ANO4, SPIC, KCTD8, TG, LRRC6, CACNA1E, KCNH1, SLC4A7, ZSCAN31, PGBD1, ZSCAN26) under selection in MAY and in ZMA, respectively. Considering each breed separately, four candidate genes (i.e. GBE1, GPC5, SLC6A2 and ZMAT4) were detected in MAY population and four (i.e. CPS1, KCND2, KCTD8 and KCNH1) in ZMA population by two *Rsb* tests.

Overall, a total of 27 and 47 candidate genes under selection were detected in at least one of the above tests for MAY and ZMA, respectively. To obtain a global view of the main gene functions under selection, these two lists of candidate genes were annotated using IPA software. The main functional categories, in which these genes are involved, are listed in Table S3 (see Tables S4 and S5 for an exhaustive list of their functional annotation). For MAY, the top five significant functions belonging to the three IPA main categories (i.e. Diseases and Disorders, Molecular and Cellular Functions and Physiological System Development and Function) were related to inflammation (‘Inflammatory Disease’), cancer and cell death (‘Cancer’ and ‘Cell Death and Survival’), gastrointestinal and hepatic diseases (‘Hepatic Disease’,’Gastrointestinal disease’), nervous system (‘Nervous System Development and Function’), organ and tissue development and morphology (‘Organismal Injury and Abnormalities’, ‘Cell Morphology’, ‘Cellular development’, ‘Embryonic Development’, ‘Organ development’, ‘Organ morphology’ and ‘Organismal development’) and inflammation (‘Inflammatory Disease’). For ZMA, the top five significant functions were related to cancer, dermatology (‘Dermatological Diseases and Conditions’), organ and embryonic development (‘Organismal Injury and Abnormalities’, ‘Organismal Development’, ‘Embryonic Development’, ‘Organ Development’), metabolism and gastrointestinal disease (‘Molecular Transport’, ‘Lipid Metabolism’, ‘Small Molecule Biochemistry’, ‘Gastrointestinal Disease’), respiratory (‘Respiratory System Development and Function’) and nervous (‘Nervous System Development and Function’) systems (Table S3). Interestingly, many functional categories or subcategories (9 among the top five significant functions) were found in both popu-lations such as functions related to cancer, nervous system, organ development and gastrointestinal disease.

Moreover, candidate genes identified in each breed participated to one significant network (Table S6) which included 85% and 93% of candidates genes for MAY (score=54) and ZMA (score=108) breeds, respectively (Figures 5a and 5b). Ten molecules are in common between both networks (i.e. AR, GRIK2, HMGCS2, LPCAT2, MAP3K10, MAP3K11, PHGDH, TRIM10, TRIM15 and kynurenic acid) among which six were detected as candidate genes under selection (i.e. GRIK2, HMGCS2, LPCAT2, PHGDH, TRIM10, TRIM15). Candidate genes were located in periphery or sub-periphery of both networks and no main node were detected under selection, except TNFSR11, associated with a significant *iHS* for ZMA breed. The functional annotation of each network gave a more complete picture of the functions involved in adaptation of MAY and ZMA breeds (Tables S7, S8 and S9). Molecules integrated in the MAY network play a role in carbohydrate metabolism, development, nervous and respiratory systems function, dermatology, inflammation and behavior. The ZMA network in mainly involved in cancer, in organismal abnormalities, cell signaling growth and proliferation, reproductive system, dermatology, gastrointestinal disease, endocrine and nervous systems and inflammation.

**Figure 5.**
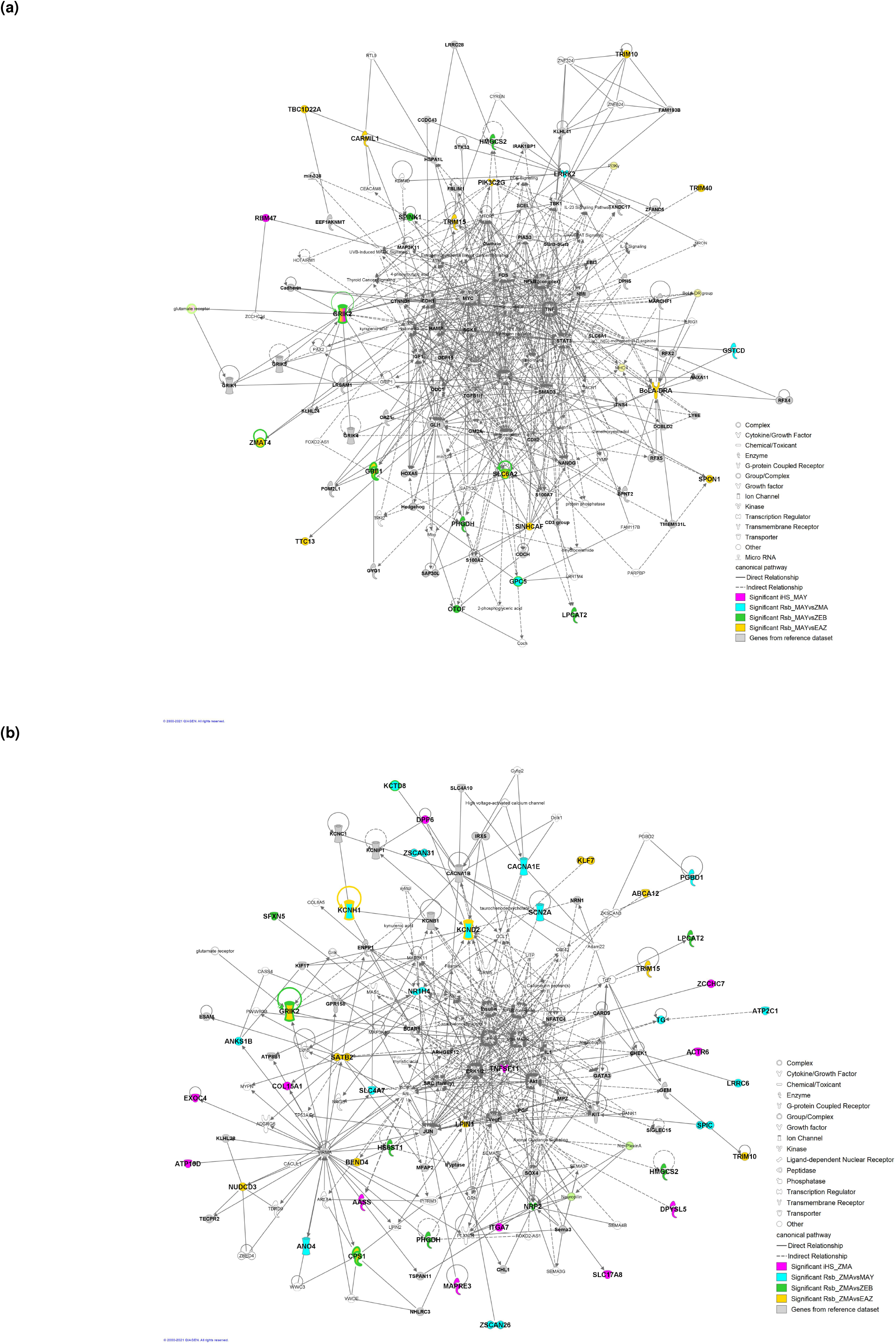
Gene networks including candidate genes under selection detected in at least one *iHS* and *Rsb* test in the genome of breeds from Mayotte (**a**) and Madagascar (**b**)

## Discussion

The genetic characterization of Zebus from Mayotte and Madagascar (MAY and ZMA), two geographically closely related island cattle populations from the Western Indian Ocean, showed that they share a similar recent and rather complex demographic history. In particular, we detected in these two populations a weak African taurine ancestry and a predominant indicine ancestry. The indicine ancestry in MAY and ZMA was even higher than in the East-African zebu breed, the African continental breed with the highest estimated indicine ancestry proportion in our dataset. This pattern had already been reported for ZMA in some previous studies (Hanotte 2002; Gautier *et al*. 2009) but still remained difficult to explain from an historical point of view. Our demographic inference, in the form of admixture graph construction, demonstrated that the higher indicine ancestry in the MAY and ZMA populations, as compared to other African continental zebus, could actually be explained by a second pulse of Indian Zebu introgression into an already admixed African taurine x Zebu population of likely East-African origin that traces back to the 12^th^ century. These results are in agreement with the recent findings in human population dynamics in the area (Brucato *et al*. 2017, 2018) based on genetic data analysis, which report that Comorian and Malagasy communities resulted from admixture between Swahili people from East-Africa and individuals from islands of Southeast Asian (Austronesian) dating from the 8^th^-12^th^ centuries (Brucato *et al*. 2018). Migrations of Austronesian populations was actually promoted by the significant development of sea borne trade, from the Arabian Peninsula and Indonesia to East Africa and Madagascar during this period (Choudhury *et al*. 2018; Brucato *et al*. 2018; Bahbahani *et al*. 2017; Brucato *et al*. 2016). These excellent navigators should have transported indicine cattle (and probably other livestock species) towards Western Indian Ocean that must have been crossed with AFZ individuals brought from African East coast by Swahili people.

Estimation of past effective population sizes for both MAY and ZMA populations showed similar trend and allowed clarifying the timing of their recent split to the 16^th^ century which coincides with the arrival of Europeans in the area and the modification of the trade network (Newitt 1983; McPherson 1984; Beaujard 2005). Some residual cattle migration may have still remained after this split with either a differential contribution of an unknown continental population to MAY and ZMA or some weak and asymmetric gene flow between them, further discriminating between these two scenarios would require additional samples from the area that may be difficult to collect (e.g., ancient DNA). Looking forward in time, after the split of MAY and ZMA, we estimated a steep increase in their ancestral populations sizes starting from the beginning of the 16^th^ century to the 17^th^ century, which was more pronounced for ZMA probably due to the far larger size of Madagascar. This observed cattle population expansion may be related to the increase of the livestock production by islands communities, in particular to meet the growing demand of European ships in fresh meat (Newitt 1983). Indeed, as the Portuguese retained control of the East African coast, the other Europeans (Dutch, English and French), who began to enter Indian Ocean, needed revictualling ports and bases that Madagascar and the Comoros islands could offer. Cattle then represented one of the main trading resources that contributed to the economic development in this area during the early 17^th^ century. From the middle of the 17^th^ century, the population sizes of both MAY and ZMA tended to stabilize suggesting a maintenance of cattle production until the beginning of the 20^th^ century, when both population sizes started to increase again but at a more moderate rate. This period actually coincides with French colonization of both islands that possibly lead to a new development in the use of local bovine resources. As expected, the characterization and partitioning of inbreeding levels into age-related HBD classes for MAY and ZMA individuals was found consistent with the inferred recent history of their corresponding population *N*_*e*_. More specifically, in both populations, the contribution to individual inbreeding levels of HBD classes of most recent origin (with rates *R*_*k*_ *<* 64, i.e., tracing back to ancestors living less than 32 generations ago) remained limited, except for a few individuals that may result from sporadic unintentional consanguineous matings. This is thus quite encouraging from a breed conservation perspective. Conversely, the highest contributing HBD classes pointed to ancestors living between 64 and 128 generations (i.e., HBD classes with rates *R*_*k*_ =128 and *R*_*k*_ =256). Assuming a 6 year generation time (see above), this corresponds to the middle of the 13^th^ century and the middle of the 17^th^ century and thus before our estimated split and expansion of the two populations. Accordingly, very similar profile of the most remote HBD classes contribution (with *R*_*k*_ ≥ 128) are observed for both MAY and ZMA individuals since these pertain to their common ancestral population.

From the 12^th^ century (i.e., ca. 150 generations ago) onward, cattle living in Comoros and Madagascar islands have likely experienced various environmental and artificial selection pressures. To identify their footprints in the MAY and ZMA breeds, we relied on EHH-based tests considering different contrasts that our detailed inference of the demographic history helped interpreting. First, it should be noticed that only a few candidate genes were found in common in the two populations when comparing the various population-specific signals. These include i) two genes (TRIM10 and TRIM15) with significant *Rsb*_*MAY*/*EAZ*_ and *Rsb*_*ZMA*/*EAZ*_ signals; and ii) five genes (HMGSC2, PHGDH, GRIK2, MIR2285BM, LPCAT2) with significant *Rsb*_*MAY*/*ZEB*_ and *Rsb*_*ZMA*/*ZEB*_ signals. Note that among the latter, GRIK2 also displayed a significant *iHS* signal in the MAY breed. These different common footprints may be considered of older origin in response to environmental (e.g., tropical climate) and human-driven constraints imposed to the common ancestral population of MAY and ZMA breeds (i.e., predating their divergence). Among these genes, TRIM10 and TRIM15 are involved in innate immune response and were already detected under selection in Muturu cattle in Africa (Tijjani *et al*. 2019). In addition, missense mutations (with respect to the Taurine assembly) within TRIM10 were found fixed in Nelore suggesting a putative role in tropical adaptation (Júnior *et al*. 2020). HMGCS2, PHGDH and LPCAT2 are related to lipid or serine metabolism (Hegardt 1999; Vilà-Brau *et al*. 2011; Reid *et al*. 2018; Morimoto *et al*. 2014), the first two being reported as associated with some fatty acid in intramuscular fat of Nelore cattle (Cesar *et al*. 2014). GRIK2 that participates to the metabolism of glutamate, an important brain neurotransmitter (Purves *et al*. 2001), is involved in tropical cattle behavior. For instance, deficiency of this gene was found associated with a significant reduction in anxiety, fear memory and the flight time temperament phenotype in tropical cattle with varying amount of indicine ancestry (Ko *et al*. 2005; Porto-Neto *et al*. 2014).

Conversely, significant signals with *Rsb*_*MAY*/*ZMA*_ based tests only, provided insights into footprints of selection of recent origin postdating the divergence of the two populations. These signals might be related to local differences between the tropical climate of Mayotte and Madagascar. In the small Mayotte island, the climate is homogeneous with a hot, humid and rainy season during the northeastern monsoon and a cooler dry season. In contrast, Madagascar island whose surface area is 15 times greater than that of Mayotte, presents a wider variety of climates being humid tropical on the East coast, dry tropical on the West coast and temperate on the Highlands. In addition, the differential evolution of husbandry practices in the two islands after the divergence of the two populations may also have contributed to leave different selection footprints. For instance, in Mayotte, Zebus have been reared in the recent past in traditional extensive systems with small herds (a few heads) with regular close contacts with breeders. Besides, in Madagascar, the sampled individuals originated from an area where breeding practices are based on larger herds (from 4 to few hundred heads) often dispersed on large pasture territory (Zafindrajaona 1991). A higher number of ZMA driven than MAY driven signals (n=18 vs. n=3) were found with this test, as expected from the twice highest *N*_*e*_ of ZMA over the period which may result in improved detection power. Among the three genes that display significant signals in the MAY breed (LRRK2, GSTCD and GPC5), GSTCD is involved in lung function and was identified as CNVR-harbored gene associated with adaptation to hypoxia in yak population (Wang *et al*. 2019). GPC5 was associated with feed efficiency in cattle (Buitenhuis *et al*. 2014; Serão *et al*. 2013). The 18 genes with significant *Rsb*_*MAY*/*ZMA*_ signals driven by ZMA breed included ANOA4, which is located in a region under selection in New World Creole cattle and in Korean Brindle Hanwoo cattle (Gautier and Naves 2011; Kim *et al*. 2018). This gene belongs to the anoctamin protein family, whose members are expressed in sweat glands and are involved in thermal sweating, one of the mechanisms of heat tolerance (Cui and Schlessinger 2015; Jian *et al*. 2014). Within this gene list, KCTD8, encodes a potassium channel, was found associated with the Temperature Humidity Index in Mediterranean cattle breeds and also identified as a heat stress responsive gene in chicken (Flori *et al*. 2019; Sun *et al*. 2015). It is also associated with milk fat percentage in buffaloes (De Camargo *et al*. 2015) and located within selection signatures detected in dairy cattle breeds (Hayes *et al*. 2008; Flori *et al*. 2009). Potassium channels are also related to the production of prolactin, a key hormone for mammary development, milk production, hair development, thermoregulation and water balance during heat stress (Underwood and Suttle 1999; Silanikove *et al*. 2000; Czarnecki *et al*. 2003; Littlejohn *et al*. 2014). KCNH1 represents another remarkable gene since it is associated with a syndrome characterized by dysmorphism and hypertrichosis in human (Kortüm *et al*. 2015) and was associated with several climatic variables in local Mediterranean cattle breeds (Flori *et al*. 2019). Finally, the thyroglobulin gene (TG), encodes a protein required for thyroid hormone synthesis and iodine storage and contains variants associated with carcass and meat quality traits in beef cattle (Hou *et al*. 2011).

The timing of the footprints underlying the other 18 and 26 candidate genes found specific to MAY (*iHS*_*MAY*_,*Rsb*_*MAY*/*EAZ*_, *Rsb*_*MAY*/*ZEB*_) or to ZMA (*iHS*_*ZMA*_,*Rsb*_*ZMA*/*EAZ*_, *Rsb*_*ZMA*/*ZEB*_) breeds is more difficult to interpret. For instance, they may either result from recent breed-specific selective constraints or older selective constraints further relaxed in one breed. In both cases, the fact that these genes were not identified with *Rsb*_*MAY*/*ZMA*_ could be due to the overall limited power of the different tests (see above). Yet, some noticeable genes showed up in the list of these breed-specific candidate genes. For instance, among the genes specifically detected in MAY, SPINK1 promotes the progression of various types of cancer and is involved in abnormal morphology of thin skin (Lin 2021). PIK3C2G, under selection in Normande and Holstein breeds (Flori *et al*. 2009), is involved in the regulation of insulin-mediated activation of the PI3K/Akt pathway in the liver that controls metabolism (Braccini *et al*. 2015), PI3K and Akt corresponding to central nodes of the MAY network (Figure 5a). TRIM40, is involved in innate immune response and was already detected under selection in Muturu cattle in Africa (Tijjani *et al*. 2019) and BOLA-DRA, which encodes the alpha subunit of the class II Major Histocompatibility Complex, expressed at the Antigen-Presenting Cells surface, which presents peptides from foreign origin to the immune system. Among the genes specific to ZMA breed, DDP6 was associated with feed efficiency (de Camargo *et al*. 2015; Buitenhuis *et al*. 2014; Serão *et al*. 2013), COL15A1, with iris hypopigmentation in Holstein Friesan cattle (Hollmann *et al*. 2017) with a possible involvement in UV-protection and NUDCD3, with milk yield and components in Swedish cattle breeds (Ghoreishifar *et al*. 2020), this latter being under positive selection in dairy cattle breeds and across Holstein, Angus, Charolais, Brahman, and N’Dama breeds (Flori *et al*. 2009; Xu *et al*. 2015).

Overall, the functional annotation of all the candidate genes identified with at least one test and of their inferred underlying networks provided some insights into the main functions involved in adaptation of cattle to Western Indian Ocean tropical conditions and breeding practices. Some genes (e.g. SPINK1 in MAY and ANO4, KCTD8, COL15A1 and KCNH1 in ZMA) are related to cancer and skin properties and may have played a role in adaptation to Mayotte and Madagascar climates by increasing tolerance to thermal stress, humidity and UV exposition. Respiratory function in which some candidate genes are involved (e.g. GSTCD in MAY) may also be related to tropical environment adaptation, breeds from warm climates presenting lower respiration rates under heat stress to maintain their body temperature. Several genes carrying footprints of selection are involved in nervous system and cattle behavior (e.g. GRIK2 in MAY and ZMA) which indicates also a physiological adaptation to the livestock system and the proximity with breeders in both islands. Development of different organs or physiological structures, gastrointestinal and hepatic system function, metabolism and hormone production are also among the main functions targeted by selection in the two breeds (e.g. HMGCS2, PHGDH, MIR2285BM and LPCAT2, detected in MAY and ZMA breeds, GPC5 detected in MAY breed and NUDCD3, TG and DDP6 detected in ZMA breed). If these signatures of selection could be related to artificial selection on conformation and production traits, they could also be linked to the adaptation to tropical climate (e.g. through thermoregulation) and to available food and water resources. At last, several candidate genes involved in inflammatory and immune response (e.g. TRIM10, TRIM15, TRIM40, BOLA-DRA) play a role in adaptation of Mayotte and Madagascar breeds suggesting that tropical pathogens responsible for e.g. as East Coast Fever, Rift Valley Fever or blackleg documented outbreaks, may have had a deep impact on the adaptive genetic diversity of MAY and ZMA breeds (De Deken *et al*. 2007; Porter *et al*. 2016; Cêtre-Sossah *et al*. 2012; Dommergues *et al*. 2015).

This study displays for the first time a genetic characterization of the local cattle breed of Mayotte and presents a deep analysis of demographic and adaptive histories of cattle breeds from Mayotte and Madagascar islands. The genetic proximity between these two populations reflects their closely tied demographic history until the 16^th^ century before the population divergence. Their demographic history mirrors the complex pattern of human migrations and trade in Western Indian Ocean islands. Their adaptive history has been probably overall conditioned by the same selective pressures and also by some differences between climates and breeding practices in the two islands. This study highlights the great originality of Zebus from Madagascar and Mayotte compared to the cattle populations from Africa mainland and their relevance in the context of global changes, which promotes specific measures of conservation especially for Mayotte breed.

## Data availability

SNP genotyping data are available in the WIDDE database (http://widde.toulouse.inra.fr/widde/) and in the Data INRAE portal (https://data.inrae.fr/).

## Acknowledgments

We would like to thank Sitty Bahyat Chamassi et Abdou Achiraffi (Laboratoire Vétérinaire d’Analyses Départemental de Mayotte, Kaweni, Mayotte) and Anlidine Mkadara, Hidachi Attoumani, Oussoufi Saindou and Kamardine Moussa (Chambre de l’Agriculture, de la Pêche et de l’Aquaculture de Mayotte, Coconi, Mayotte) for their technical assistance and all the breeders of the Mayotte cattle breed who participated to this study. Tom Druet is Senior Research Associate from the F.R.S.-FNRS.

---

**Figure S1.**
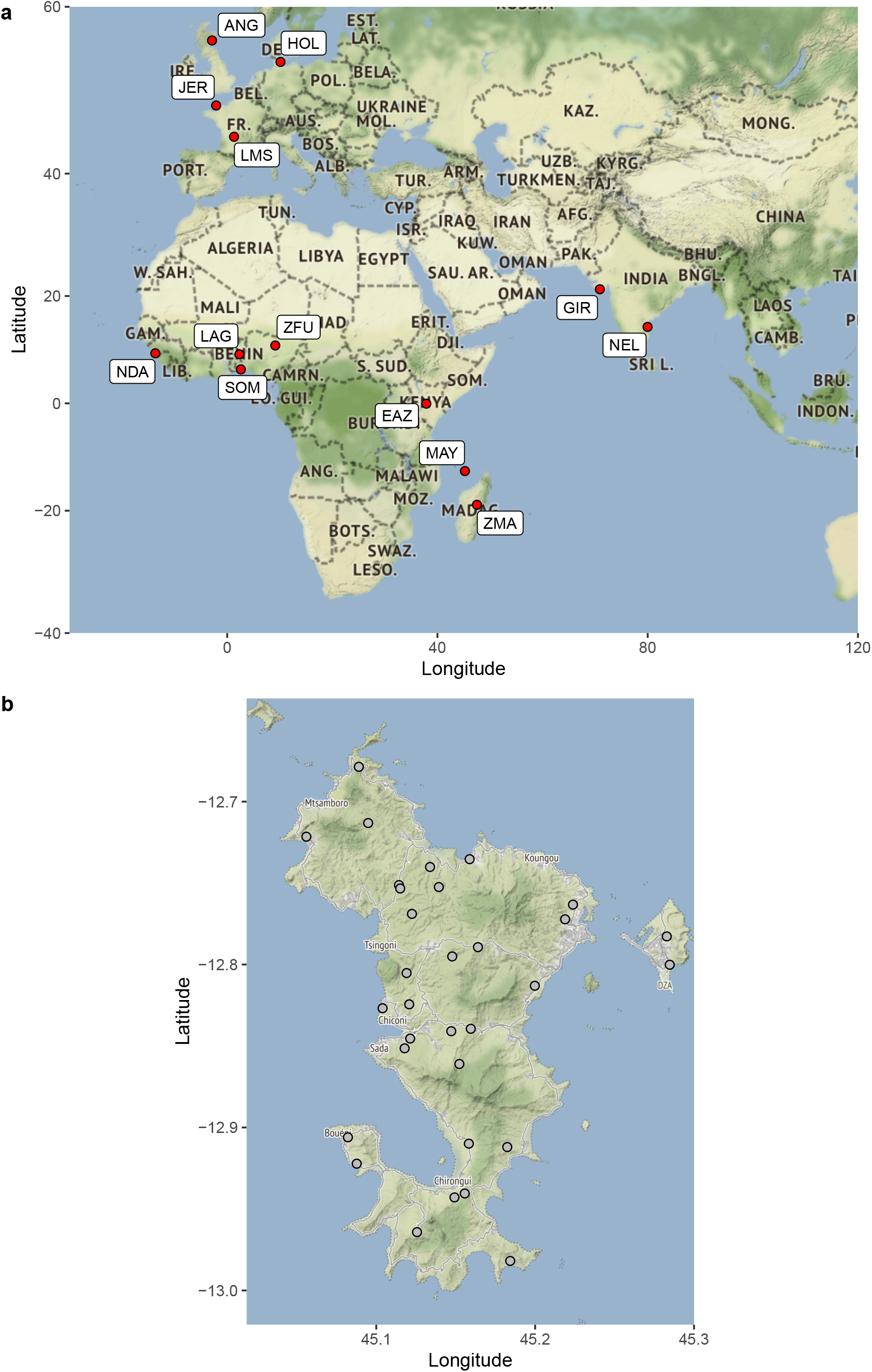
Maps and geographic coordinates of the populations studied (**a**) and the farms (**b**) where Zebus from Mayotte were sampled. Population coordinates correspond to the area of sampling except for those of Gir (GIR) and Nelore (NEL) populations which are located in their zone of origin.

**Figure S2.**
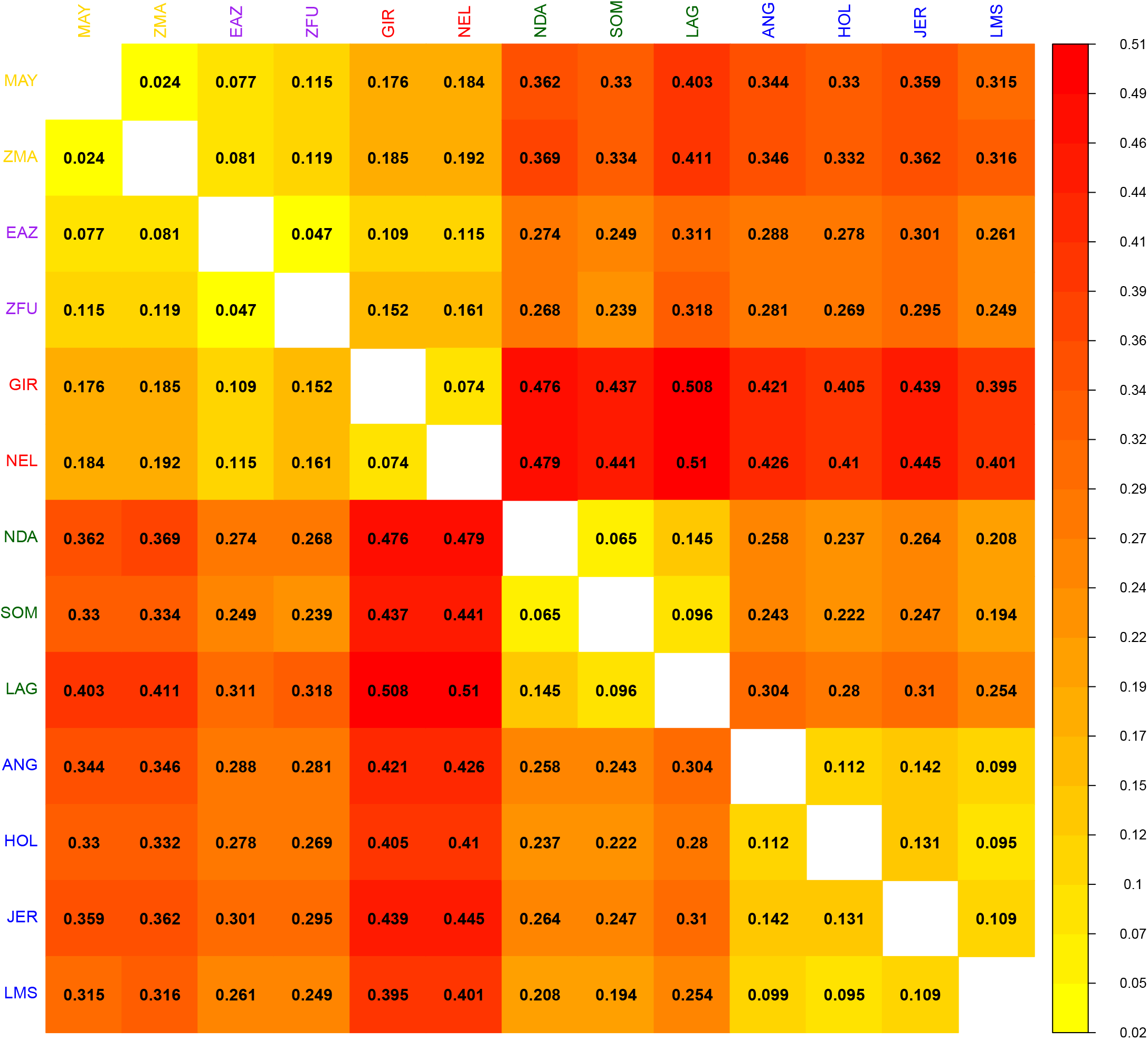
Pairwise-population *F*_*ST*_ among the all the 13 populations.

**Figure S3.**
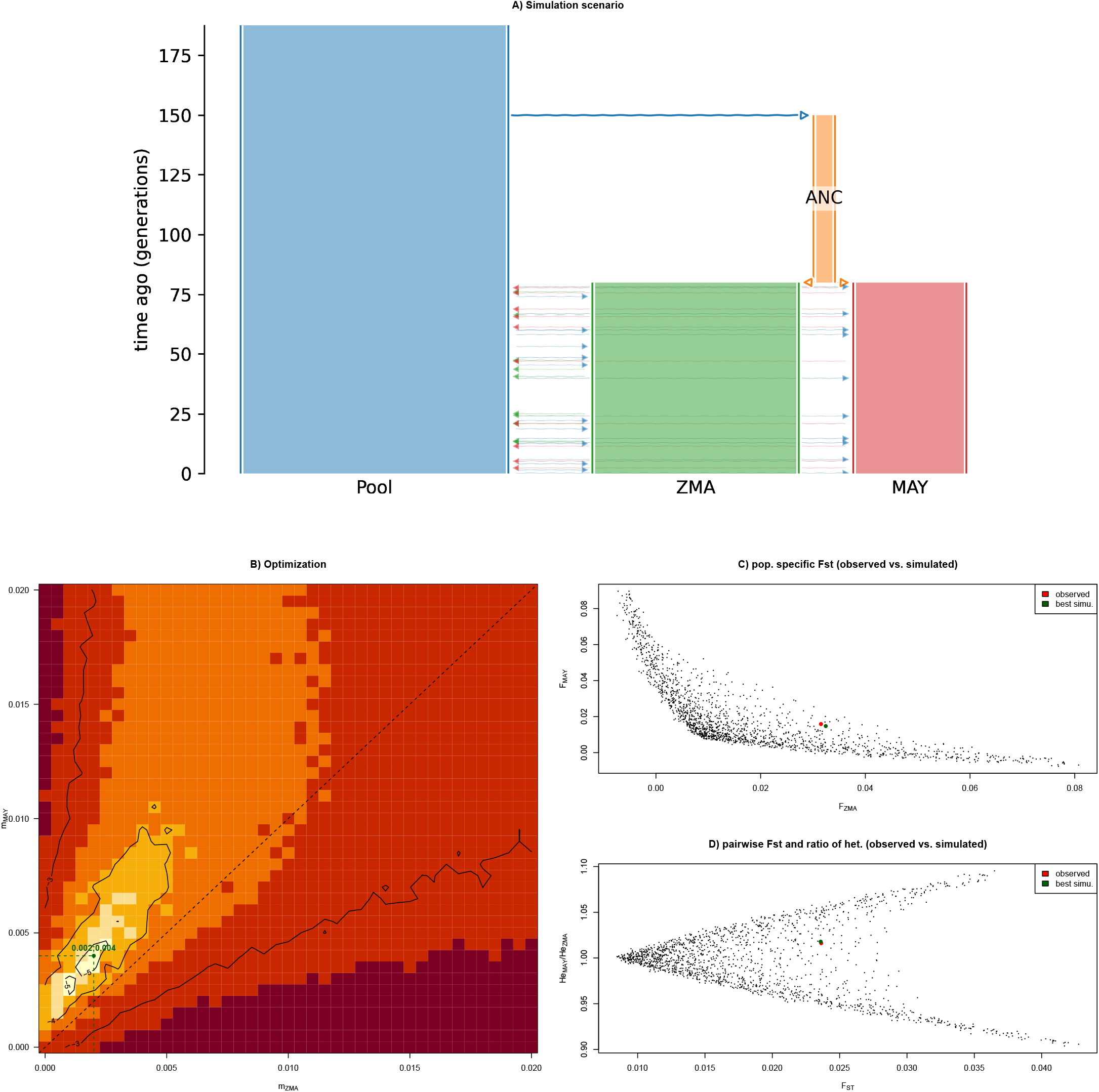
Simulation based evaluation of isolation models for ZMA and MAY with asymmetrical migration rates using msprime (Kelleher *et al*. 2016). A) Simulation scenario. Modeling of asymmetrical migration rate is performed via a (ghost) population (‘Pool’) with constant *N*_*e*_ =2500 that contribute to ZMA and MAY population with different symmetric migration rate (*m*_*ZMA*_ and *m*_*MAY*_ respectively) after the split from a common ancestral population named ‘ANC’ occurring *t* = 80 generations ago. The *N*_*e*_ for ZMA and MAY was set to 2,163 and 1,045 respectively which corresponded to the harmonic mean of the corresponding historical *N*_*e*_ estimates (Figure 3). Similarly the *N*_*e*_ for ANC was set to 250 and its divergence from the trunk population occurred *t* = 150 generations ago (i.e., the estimated timing of admixture for population ancestral to ZMA and MAY, named Ind. Oc. Zebus in Figure 2). B) Estimated distance *δ* (in a log_10_ scaled color gradient) between the observed data and data simulated over a grid of *m*_*zma*_ and *m*_*may*_ values (41 values ranging from 0 to 0.02 with a step of 0.005 for each parameter leading to a total of 1, 681 = 41 × 41 simulated data sets). Each simulated data set consisted of 250 independent recombining segments of 5 Mb (assuming per-generation and per base mutation and recombination rates equal to *µ* = *r* = 10^−8^) for 23 ZMA and 30 MAY individuals and with a 4 generations sampling decay (as for the original data). The distance simulated and observed data was computed as 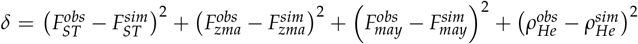 where *F* (resp. *F* is the population-specific differentiation (e.g., 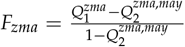) computed from poolfstat estimates of *Q* and *Q* probability of identity (i.e., 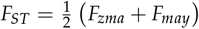) and *ρ* is the ratio of within population MAY and ZMA heterozygosities (= 1 − *Q*_1_). SNPs with a MAF*<* 0.01 (computed over MAY and ZMA combined samples) were discarded in both the observed and simulated to compute the statistics (leading to 518,315 SNPs for the observed data set and from 381,197 to 474,633 for the simulated ones). The optimal parameters values that minimized *δ* (*m*_*zma*_ = 0.002 and *m*_*may*_ = 0.004) are highlighted in green. C) Estimated *F*_*zma*_ and *F*_*may*_ for all the simulated data sets. The optimal parameters values (see B)) and the values for the observed data set are highlighted in green and red respectively. D) Same as C) with estimated pairwise *F*_*ST*_ and ratio of heterozygosities *ρ*_*He*_.

**Figure S4.**
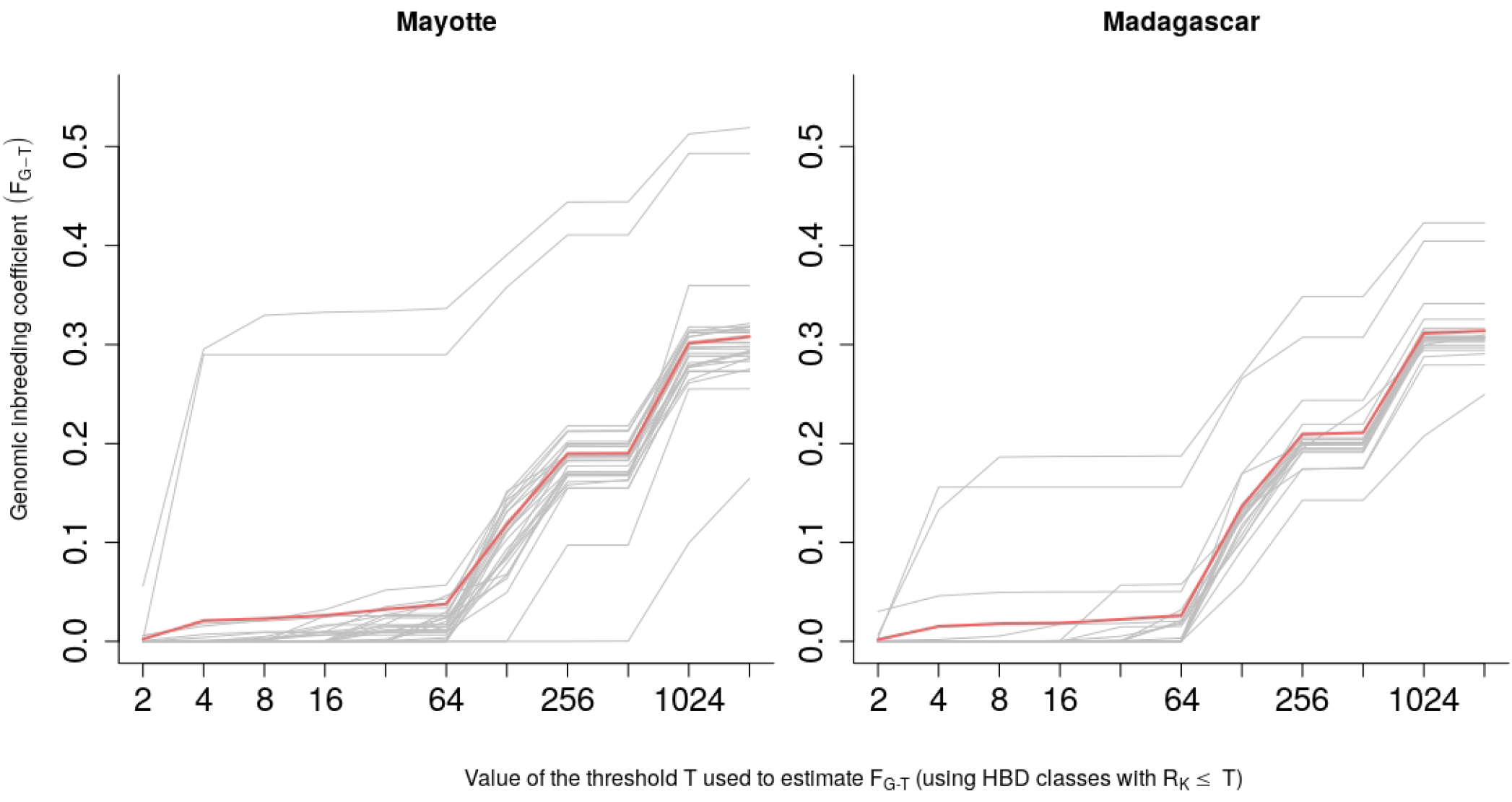
Proportion of the genome associated with different HBD classes at an individual level for Zebu from MayFreprootte (left) and Zebu from Madagascar (right). The figure was created with RzooRoh package (Bertrand et al 2019).

**Figure S5.**
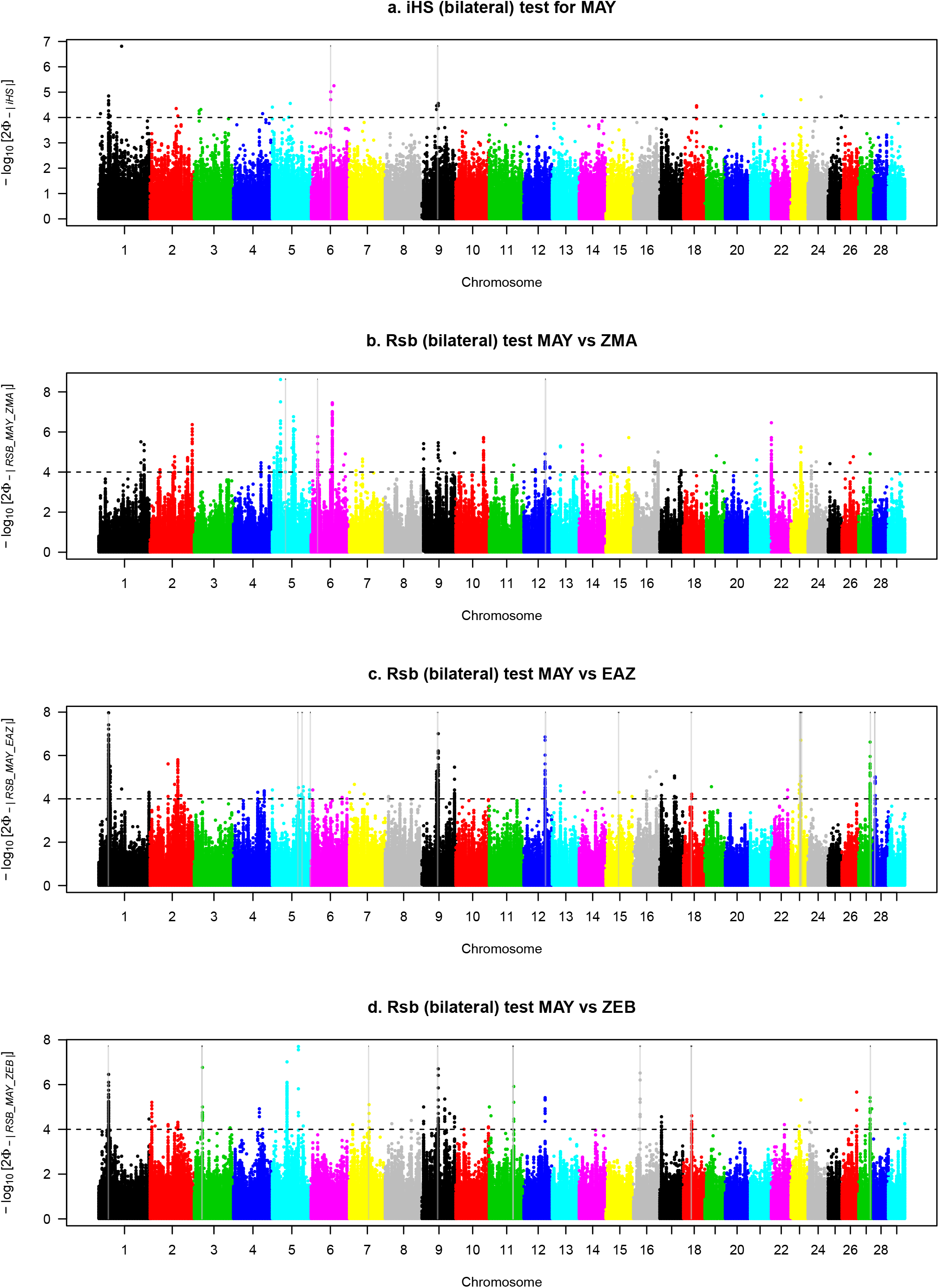
Manhattan plot of pvalues for *iHS*_*MAY*_, *Rsb*_*MAYvsZMA*_, *Rsb*_*MAYvsEAZ*_, *Rsb*_*MAYvsZEB*_ analyses. Positions of candidate genes are indicated by gray lines.

**Figure S6.**
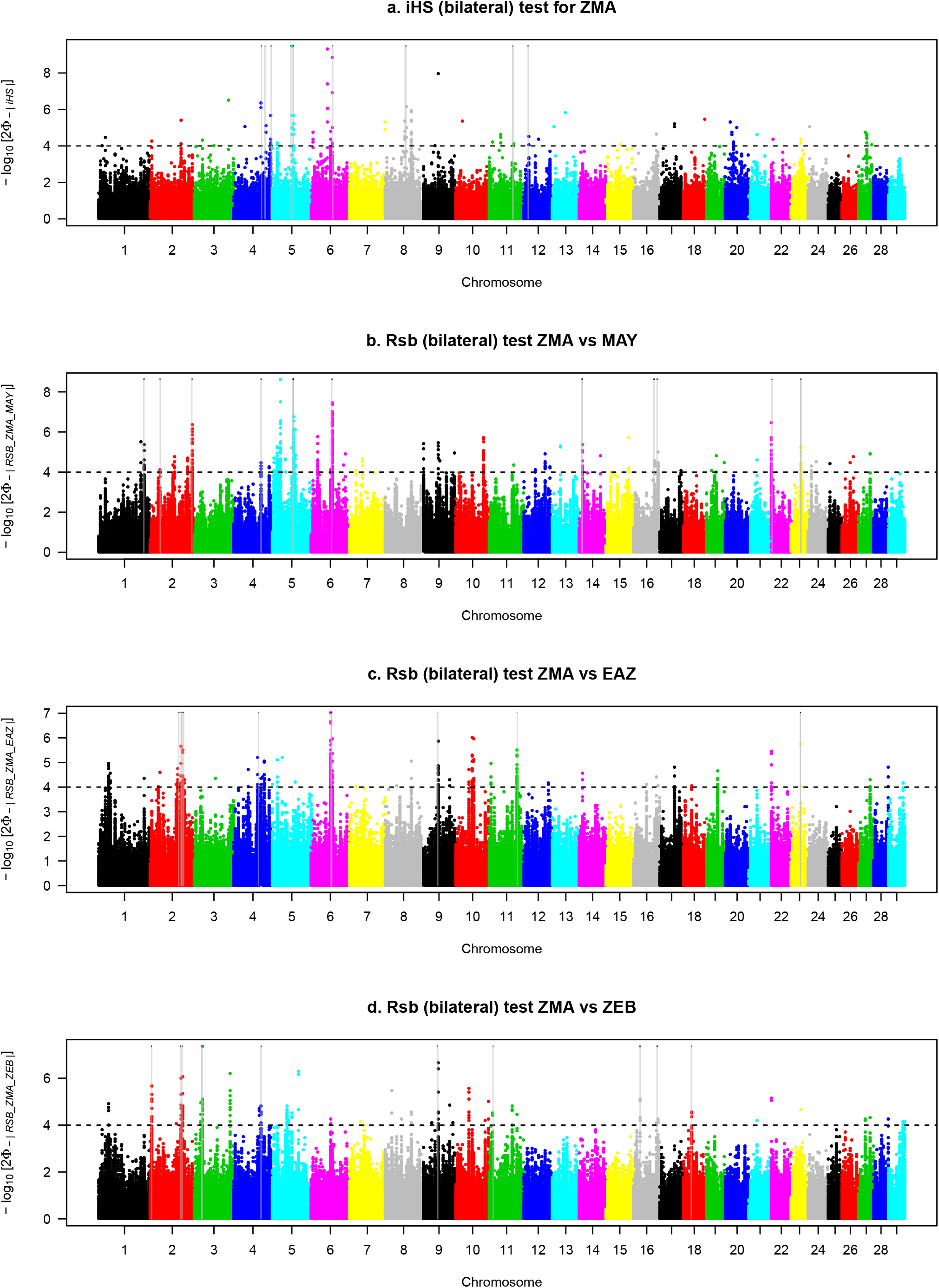
Manhattan plot of pvalues for *iHS*_*ZMA*_, *Rsb*_*ZMAvsMAY*_, *Rsb*_*ZMAvsEAZ*_, *Rsb*_*ZMAvsZEB*_ analyses. Positions of candidate genes are indicated by gray lines.

## Literature cited

Alexander DH, Novembre J, Lange K. 2009. Fast model-based estimation of ancestry in unrelated individuals. Genome Research. 19:1655–1664.

Bahbahani H, Tijjani A, Mukasa C, Wragg D, Almathen F, Nash O, Akpa GN, Mbole-Kariuki M, Malla S, Woolhouse M et al. 2017. Signatures of Selection for Environmental Adaptation and Zebu x Taurine Hybrid Fitness in East African Shorthorn Zebu. Frontiers in Genetics. 8.

Beaujard P. 2005. The Indian Ocean in Eurasian and African world-systems before the sixteenth century. Journal of World History. 16:411–+. WOS:000235458400002.

Beaujard P. 2007. East Africa, the Comoros Islands and Madagascar before the sixteenth century: On a neglected part of the world system. Azania: Archaeological Research in Africa. 42:15–35.

Beaujard P. 2011. The first migrants to Madagascar and their introduction of plants: linguistic and ethnological evidence. Azania-Archaeological Research in Africa. 46:169–189. WOS:000293472700004.

Beaujard P. 2015. East Africa and oceanic exchange networks between the first and the fifteenth centuries. Afriques-Debats Methodes Et Terrains D Histoire. 6. WOS:000420334300001.

Beaujard P. 2019a. The Worlds of the Indian Ocean: A Global History: Volume 1: From the Fourth Millennium BCE to the Sixth Century CE. volume 1. Cambridge University Press. Cambridge.

Beaujard P. 2019b. The worlds of the Indian Ocean: A global history: Volume 2 : From the Seventh Century to the Fifteenth Century CE. volume 2. Cambridge University Press. Cambridge, United Kingdom; New York, NY.

Bertrand AR, Kadri NK, Flori L, Gautier M, Druet T. 2019. Rzooroh: an r package to characterize individual genomic autozygosity and identify homozygous-by-descent segments. Methods in Ecology and Evolution. 10:860–866.

Bock R, Kingston T, De Vos A. 1999. Effect of breed of cattle on transmission rate and innate resistance to infection with babesia bovis and b bigemina transmitted by boophilus microplus.

Boivin N, Crowther A, Helm R, Fuller DQ. 2013. East Africa and Madagascar in the Indian Ocean world. Journal of World Prehistory. 26:213–281.

Braccini L, Ciraolo E, Campa CC, Perino A, Longo DL, Tibolla G, Pregnolato M, Cao Y, Tassone B, Damilano F et al. 2015. PI3K-C2Y is a Rab5 effector selectively controlling endosomal Akt2 activation downstream of insulin signalling. Nature Communications. 6:1–15.

Brucato N, Fernandes V, Mazières S, Kusuma P, Cox MP, Ng’ang’a JW, Omar M, Simeone-Senelle MC, Frassati C, Alshamali F et al. 2018. The Comoros Show the Earliest Austronesian Gene Flow into the Swahili Corridor. The American Journal of Human Genetics. 102:58–68.

Brucato N, Kusuma P, Beaujard P, Sudoyo H, Cox MP, Ricaut FX. 2017. Genomic admixture tracks pulses of economic activity over 2,000 years in the Indian Ocean trading network. Scientific Reports. 7:2919.

Brucato N, Kusuma P, Cox MP, Pierron D, Purnomo GA, Adelaar A, Kivisild T, Letellier T, Sudoyo H, Ricaut FX. 2016. Malagasy Genetic Ancestry Comes from an Historical Malay Trading Post in Southeast Borneo. Molecular Biology and Evolution. 33:2396–2400.

Buitenhuis B, Janss LL, Poulsen NA, Larsen LB, Larsen MK, Sørensen P. 2014. Genome-wide association and biological pathway analysis for milk-fat composition in danish holstein and danish jersey cattle. BMC genomics. 15:1–11.

Cesar AS, Regitano LC, Mourão GB, Tullio RR, Lanna DP, Nassu RT, Mudado MA, Oliveira PS, do Nascimento ML, Chaves AS et al. 2014. Genome-wide association study for intramuscular fat deposition and composition in nellore cattle. BMC genetics. 15:1–15.

Cheke A. 2010. The timing of arrival of humans and their commensal animals on Western Indian Ocean oceanic islands. Phelsuma. 18:38–69.

Choudhury A, Aron S, Sengupta D, Hazelhurst S, Ramsay M. 2018. African genetic diversity provides novel insights into evolutionary history and local adaptations. Human Molecular Genetics. 27:R209–R218.

Cui CY, Schlessinger D. 2015. Eccrine sweat gland development and sweat secretion. Experimental Dermatology. 24:644–650.

Czarnecki A, Dufy-Barbe L, Huet S, Odessa MF, Bresson-Bepoldin L. 2003. Potassium channel expression level is dependent on the proliferation state in the GH3 pituitary cell line. American Journal of Physiology-Cell Physiology. 284:C1054–C1064.

Cêtre-Sossah C, Pédarrieu A, Guis H, Defernez C, Bouloy M, Favre J, Girard S, Cardinale E, Albina E. 2012. Prevalence of Rift Valley Fever among Ruminants, Mayotte. Emerging Infectious Diseases. 18:972–975.

De Camargo G, Aspilcueta-Borquis RR, Fortes M, Porto-Neto R, Cardoso DF, Santos D, Lehnert S, Reverter A, Moore S, Tonhati H. 2015. Prospecting major genes in dairy buffaloes. BMC genomics. 16:1–14.

de Camargo GMF, Porto-Neto LR, Kelly MJ, Bunch RJ, McWilliam SM, Tonhati H, Lehnert SA, Fortes MR, Moore SS. 2015. Non-synonymous mutations mapped to chromosome x associated with andrological and growth traits in beef cattle. BMC genomics. 16:1–10.

De Deken R, Martin V, Saido A, Madder M, Brandt J, Geysen D. 2007. An outbreak of East Coast Fever on the Comoros: A consequence of the import of immunised cattle from Tanzania? Veterinary Parasitology. 143:245–253.

Dommergues L, Pannequin M, Cavalerie L, Cardinale E. 2015. Etude épidémiologique sur le charbon symptomatique à Mayotte (2014). Technical report. Coopadem - GDS Mayotte.

Druet T, Gautier M. 2017. A model-based approach to characterize individual inbreeding at both global and local genomic scales. Molecular Ecology. 26:5820–5841.

Druet T, Gautier M. 2021. An improved hidden markov model for the characterization of homozygous-by-descent segments in individual genomes. bioRxiv..

Flori L, Fritz S, Jaffrézic F, Boussaha M, Gut I, Heath S, Foulley JL, Gautier M. 2009. The genome response to artificial selection: a case study in dairy cattle. PloS One. 4:e6595.

Flori L, Moazami-Goudarzi K, Alary V, Araba A, Boujenane I, Boushaba N, Casabianca F, Casu S, Ciampolini R, D’Acier AC et al. 2019. A genomic map of climate adaptation in Mediter-ranean cattle breeds. Molecular Ecology. 28:1009–1029.

Flori L, Thevenon S, Dayo GK, Senou M, Sylla S, Berthier D, Moazami-Goudarzi K, Gautier M. 2014. Adaptive admixture in the West African bovine hybrid zone: insight from the Borgou population. Molecular Ecology. 23:3241–3257. WOS:000338014900009.

France M. 2011. Synthèse illustrée du recensement agricole 2010. Technical report. Ministère de l’Agriculture, de l’Alimentation, de la Pêche, de la Ruralité et de l’Aménagement du Territoire. Mayotte.

Fuller D, Boivin N, Hoogervorst T, Allaby R. 2011. Across the Indian Ocean: the prehistoric movement of plants and animals. Antiquity. 85:544–558.

Fuller DQ, Boivin N. 2009. Crops, cattle and commensals across the Indian Ocean. Etudes Ocean Indien. pp. 13–46.

Gautier M, Faraut T, Moazami-Goudarzi K, Navratil V, Foglio M, Grohs C, Boland A, Garnier JG, Boichard D, Lathrop GM et al. 2007. Genetic and Haplotypic Structure in 14 European and African Cattle Breeds. Genetics. 177:1059–1070.

Gautier M, Flori L, Riebler A, Jaffrézic F, Laloé D, Gut I, Moazami-Goudarzi K, Foulley JL. 2009. A whole genome Bayesian scan for adaptive genetic divergence in West African cattle. BMC genomics. 10:550.

Gautier M, Klassmann A, Vitalis R. 2017. rehh 2.0 : a reimplementation of the R package rehh to detect positive selection from haplotype structure. Molecular Ecology Resources. 17:78–90.

Gautier M, Laloë D, Moazami-Goudarzi K. 2010. Insights into the genetic history of French cattle from dense SNP data on 47 worldwide breeds. PloS One. 5.

Gautier M, Naves M. 2011. Footprints of selection in the ancestral admixture of a New World Creole cattle breed. Molecular Ecology. 20:3128–3143.

Gautier M, Vitalis R, Flori L, Estoup A. 2021. f-statistics estimation and admixture graph construction with Pool-Seq or allele count data using the R package poolfstat. bioRxiv. 2021.05.28.445945.

Ghoreishifar SM, Eriksson S, Johansson AM, Khansefid M, Moghaddaszadeh-Ahrabi S, Parna N, Davoudi P, Javanmard A. 2020. Signatures of selection reveal candidate genes involved in economic traits and cold acclimation in five swedish cattle breeds. Genetics Selection Evolution. 52:1–15.

Glass EJ, Preston PM, Springbett A, Craigmile S, Kirvar E, Wilkie G, Brown CD. 2005. Bos taurus and bos indicus (sahiwal) calves respond differently to infection with theileria annulata and produce markedly different levels of acute phase proteins. International journal for parasitology. 35:337–347.

Hanotte O. 2002. African Pastoralism: Genetic Imprints of Origins and Migrations. Science. 296:336–339.

Hansen P. 2004. Physiological and cellular adaptations of zebu cattle to thermal stress. Animal reproduction science. 82:349–360.

Hayes BJ, Lien S, Nilsen H, Olsen HG, Berg P, Maceachern S, Potter S, Meuwissen THE. 2008. The origin of selection signatures on bovine chromosome 6. Animal Genetics. 39:105–111.

Hegardt FG. 1999. Mitochondrial 3-hydroxy-3-methylglutaryl-coa synthase: a control enzyme in ketogenesis. Biochemical Journal. 338:569–582.

Hollmann AK, Bleyer M, Tipold A, Neßler JN, Wemheuer WE, Schütz E, Brenig B. 2017. A genome-wide association study reveals a locus for bilateral iridal hypopigmentation in holstein friesian cattle. BMC genetics. 18:1–8.

Hou GY, Yuan ZR, Zhou HL, Zhang LP, Li JY, Gao X, Wang DJ, Gao HJ, Xu SZ. 2011. Association of thyroglobulin gene variants with carcass and meat quality traits in beef cattle. Molecular biology reports. 38:4705–4708.

Jian W, Duangjinda M, Vajrabukka C, Katawatin S. 2014. Differences of skin morphology in Bos indicus, Bos taurus, and their crossbreds. International Journal of Biometeorology. 58:1087–1094.

Júnior GAF, de Oliveira HN, Carvalheiro R, Cardoso DF, Fonseca LFS, Ventura RV, de Albuquerque LG. 2020. Whole-genome sequencing provides new insights into genetic mechanisms of tropical adaptation in nellore (bos primigenius indicus). Scientific reports. 10:1–7.

Kadri NK, Harland C, Faux P, Cambisano N, Karim L, Coppieters W, Fritz S, Mullaart E, Baurain D, Boichard D et al. 2016. Coding and noncoding variants in hfm1, mlh3, msh4, msh5, rnf212, and rnf212b affect recombination rate in cattle. Genome Research..

Keightley PD, Eyre-Walker A. 2000. Deleterious mutations and the evolution of sex. Science. 290:331–3.

Kelleher J, Etheridge AM, McVean G. 2016. Efficient coalescent simulation and genealogical analysis for large sample sizes. PLoS Computational Biology. 12:e1004842.

Kim S, Cheong HS, Shin HD, Lee SS, Roh HJ, Jeon DY, Cho CY. 2018. Genetic diversity and divergence among Korean cattle breeds assessed using a BovineHD single-nucleotide polymorphism chip. Asian-Australasian Journal of Animal Sciences. 31:1691–1699.

Klassmann A, Gautier M. 2020. Detecting selection using extended haplotype homozygosity-based statistics on unphased or unpolarized data. Authorea. 10.22541/au.160405572.29972398/v1.

Ko S, Zhao MG, Toyoda H, Qiu CS, Zhuo M. 2005. Altered Behavioral Responses to Noxious Stimuli and Fear in Glutamate Receptor 5 (GluR5)-or GluR6-Deficient Mice. Journal of Neuroscience. 25:977–984.

Kortüm F, Caputo V, Bauer CK, Stella L, Ciolfi A, Alawi M, Bocchinfuso G, Flex E, Paolacci S, Dentici ML et al. 2015. Mutations in kcnh1 and atp6v1b2 cause zimmermann-laband syndrome. Nature genetics. 47:661–667.

Lin TC. 2021. Functional roles of spink1 in cancers. International Journal of Molecular Sciences. 22:3814.

Lipson M. 2020. Applying f-statistics and admixture graphs: Theory and examples. Molecular Ecology Resources. 20:1658–1667.

Littlejohn MD, Henty KM, Tiplady K, Johnson T, Harland C, Lopdell T, Sherlock RG, Li W, Lukefahr SD, Shanks BC et al. 2014. Functionally reciprocal mutations of the prolactin signalling pathway define hairy and slick cattle. Nature Communications. 5:1–8.

Loh PR, Lipson M, Patterson N, Moorjani P, Pickrell JK, Reich D, Berger B. 2013. Inferring Admixture Histories of Human Populations Using Linkage Disequilibrium. Genetics. 193:1233–1254.

Mattioli R, Bah M, Kora S, Cassama M, Clifford D. 1995. Susceptibility to different tick genera in gambian n’dama and gobra zebu cattle exposed to naturally occurring tick infestations. Tropical animal health and production. 27:95–105.

McPherson K. 1984. Cultural exchange in the indian ocean region. Westerly. 29:5–16.

Ministère de l’Agriculture dledlpdM. 2007. Recensement de l’agriculture (RA) Campagne agricole 2004-2005 - Tome 1 Généralités, méthodologies et principaux résultats. Technical report. Ministère de l’Agriculture, de l’élevage et de la pêche de Madagascar.

Morimoto R, Shindou H, Tarui M, Shimizu T. 2014. Rapid production of platelet-activating factor is induced by protein kinase cα-mediated phosphorylation of lysophosphatidylcholine acyltransferase 2 protein. Journal of Biological Chemistry. 289:15566–15576.

Newitt M. 1983. The comoro islands in indian ocean trade before the 19th century (les comores et le commerce dans l’océan indien avant le xixe siècle). Cahiers d’études africaines. pp. 139–165.

Ouvrard M, Magnier J, Raoul S, Anlidine M, Attoumani H, Oussoufi A, Kamardine M, Janelle J, Naves M, Flori L et al. 2018. Caractérisation phénotypique et génétique : le cas du zébu mahorais.

Patterson N, Price AL, Reich D. 2006. Population Structure and Eigenanalysis. PLOS Genetics. 2:e190.

Patterson NJ, Moorjani P, Luo Y, Mallick S, Rohland N, Zhan Y, Genschoreck T, Webster T, Reich D. 2012. Ancient Admixture in Human History. Genetics. p. genetics.112.145037.

Pauly M. 2013. Acoua-Agnala M’kiri, Mayotte (976), archéologie d’une localité médiévale (XIe-XVe siècles), entre Afrique et Madagascar. NYAME AKUMA. pp. 73–90.

Pickrell JK, Patterson N, Loh PR, Lipson M, Berger B, Stoneking M, Pakendorf B, Reich D. 2014. Ancient west Eurasian ancestry in southern and eastern Africa. Proceedings of the National Academy of Sciences. 111:2632–2637.

Porter V. 2007. Cattle : A Handbook to the Breeds of the World. Crowood.

Porter V, Alderson L, Hall SJG, Sponenberg DP. 2016. Mason’s World Encyclopedia of Livestock Breeds and Breeding, 2 Volume Pack. CABI.

Porto-Neto LR, Reverter A, Prayaga KC, Chan EK, Johnston DJ, Hawken RJ, Fordyce G, Garcia JF, Sonstegard TS, Bolormaa S et al. 2014. The genetic architecture of climatic adaptation of tropical cattle. Plos one. 9:e113284.

Purves D, Augustine GJ, Fitzpatrick D, Katz LC, Lamantia AS, McNamara JO, Williams SM. 2001. Neuroscience. Sinauer Associates. second edition.

Reid MA, Allen AE, Liu S, Liberti MV, Liu P, Liu X, Dai Z, Gao X, Wang Q, Liu Y et al. 2018. Serine synthesis through phgdh coordinates nucleotide levels by maintaining central carbon metabolism. Nature communications. 9:1–11.

Rosen BD, Bickhart DM, Schnabel RD, Koren S, Elsik CG, Tseng E, Rowan TN, Low WY, Zimin A, Couldrey C et al. 2020. De novo assembly of the cattle reference genome with single-molecule sequencing. GigaScience. 9. giaa021.

Santiago E, Novo I, Pardiñas AF, Saura M, Wang J, Caballero A. 2020. Recent Demographic History Inferred by High-Resolution Analysis of Linkage Disequilibrium. Molecular Biology and Evolution. 37:3642–3653.

Scheet P, Stephens M. 2006. A Fast and Flexible Statistical Model for Large-Scale Population Genotype Data: Applications to Inferring Missing Genotypes and Haplotypic Phase. American Journal of Human Genetics. 78:629–644.

Sempéré G, Moazami-Goudarzi K, Eggen A, Laloë D, Gautier M, Flori L. 2015. WIDDE : a Web-Interfaced next generation database for genetic diversity exploration, with a first application in cattle. BMC Genomics. 16.

Serão NV, González-Peña D, Beever JE, Faulkner DB, Southey BR, Rodriguez-Zas SL. 2013. Single nucleotide polymorphisms and haplotypes associated with feed efficiency in beef cattle. BMC genetics. 14:1–20.

Silanikove N, Shamay A, Shinder D, Moran A. 2000. Stress down regulates milk yield in cows by plasmin induced B-casein product that blocks K+ channels on the apical membranes. Life Sciences. 67:2201–2212.

Sun L, Lamont SJ, Cooksey AM, McCarthy F, Tudor CO, Vijay-Shanker K, DeRita RM, Rothschild M, Ashwell C, Persia ME et al. 2015. Transcriptome response to heat stress in a chicken hepatocellular carcinoma cell line. Cell Stress and Chaperones. 20:939–950.

Tang K, Thornton KR, Stoneking M. 2007. A New Approach for Using Genome Scans to Detect Recent Positive Selection in the Human Genome. PLoS Biology. 5:e171.

Tijjani A, Utsunomiya YT, Ezekwe AG, Nashiru O, Hanotte O. 2019. Genome sequence analysis reveals selection signatures in endangered trypanotolerant west african muturu cattle. Frontiers in genetics. 10:442.

Underwood E, Suttle N. 1999. The mineral nutrition of livestock 3rd edition. CABI.

Vilà-Brau A, De Sousa-Coelho AL, Mayordomo C, Haro D, Marrero PF. 2011. Human hmgcs2 regulates mitochondrial fatty acid oxidation and fgf21 expression in hepg2 cell line. Journal of Biological Chemistry. 286:20423–20430.

Voight BF, Kudaravalli S, Wen X, Pritchard JK. 2006. A Map of Recent Positive Selection in the Human Genome. PLOS Biology. 4:e72.

Wang H, Chai Z, Hu D, Ji Q, Xin J, Zhang C, Zhong J. 2019. A global analysis of cnvs in diverse yak populations using whole-genome resequencing. BMC genomics. 20:1–12.

Wang M, Kong L. 2019. pblat: a multithread blat algorithm speeding up aligning sequences to genomes. BMC Bioinformatics. 20:28.

Wickham H. 2016. ggplot2 : Elegant Graphics for Data Analysis. Springer-Verlag New York..

Xu L, Bickhart DM, Cole JB, Schroeder SG, Song J, Tassell CPV, Sonstegard TS, Liu GE. 2015. Genomic signatures reveal new evidences for selection of important traits in domestic cattle. Molecular biology and evolution. 32:711–725.

Zafindrajaona PS. 1991. Profils genetiques du zebu malgache. thesis. Paris 11.

Zafindrajaona PS, Lauvergne J. 1993. Comparaison de populations de zébu malgache à l’aide des distances génétiques. Genetics, Selection, Evolution : GSE. 25:373–395.

